# Dicer1 promotes Aβ clearance via blocking B2 RNA-mediated repression of apolipoprotein E

**DOI:** 10.1101/2020.08.20.258749

**Authors:** Yan Wang, Meiling Lian, Liping Song, Shengzhou Wu

## Abstract

Metabolism of β-amyloid is critical for healthy brain. Decreased clearance of β-amyloid is associated with ensued accumulation of amyloid peptide, culminating in formation of senile plaques, a neuropathological hallmark of Alzheimer’s disease(AD). Apolipoprotein E (*APOE*), a lipoprotein for phospholipid and cholesterol metabolism, is predominantly synthesized by glia in the central nervous system, controlling Aβ aggregation and metabolism. By use of stereotactic injection and a Morris water maze, we found that delivery of Dicer1-expressing adenovirus into the hippocampus of an animal model of AD mice APPswe/PSEN1deltaE9 significantly improved spatial memory. The effect was associated with reduced amyloid peptides in the hippocampus which were analyzed with immunofluorescence and enzyme-linked immunosorbent assay. With western blot, quantitative real-time PCR, fluorescence *in situ* hybridization, and northern blot, Dicer1 overexpression increased apolipoprotein E (APOE) and concomitantly decreased B2 RNA in the hippocampus of the AD mice and in astrocyte cultures whereas transfection of B2 Mm2 RNA decreased *APOE* mRNA and protein levels in astrocyte cultures. Further, human or mouse *APOE* mRNA was found containing Alu RNA or its equivalent, B2 Mm2 RNA, locating downstream of its 3’-untranslated region (UTR), respectively. The 3’-UTR or 3’-UTR in conjunction with the downstream Alu/B2 RNA were cloned into a luciferase reporter; with dual-luciferase assay, we found that simultaneous transfection of Dicer1 siRNA or Alu/B2 RNA decreased the corresponding luciferase activities which suggest that Alu RNA mediated *APOE* mRNA degradation. Altogether, Dicer1 expression mediated amyloid peptide clearance by increasing APOE via blocking B2 RNA-mediated *APOE* mRNA degradation.

## 1. Introduction

The neuropathology hallmarks of Alzheimer’s disease (AD) include extracellular deposition of senile plaques, intracellular deposition of neurofibrillary tangles, and diffusible loss of synapses and neurons. Amyloid peptide, the major component of amyloid plaque, induces synapse loss, oxidative stress, calcium dyshomeostasis, and insulin resistance[1–4]. Thus, elimination of amyloid peptide from the brain such as β-amyloid immunotherapy has been regarded as a promising strategy to modify the progression of AD. Unfortunately, phase III clinical trial with solanezumab has failed to improve cognitive function in AD patients[5], which suggests that targeting multiple pathogenic factors is vital for treating AD.

Apolipoprotein E (*APOE*) polymorphism is the strongest risk factor for late-onset AD. The underlying mechanism of APOE involved in AD is possibly due to its effects on Aβ metabolism and clearance. In AD, Aβ production is not different, but clearance of Aβ40 or Aβ42 is decreased, compared with age-matched adults[6]. Aβ clearance is largely dependent on APOE in that APOE promotes soluble Aβ binding, internalization and subsequent degradation by glia, through low-density lipoprotein receptor-related protein 1 and heparan sulfate proteoglycan[7]. Compared with *APOE* allelic variants *ε2* and *APOE* ε3, APOE\** ε4 is less efficient to facilitate Aβ cellular uptake and degradation[8] and is also inefficient to drive Aβ efflux through blood-brain barrier. In the central nervous system, APOE is mainly produced by astrocytes and microglia[9] and is lipidated by ATP-binding cassette transporter A1 or G1 (ABCA1/ABCG1) which delivers cholesterol to APOE, without which clearance of Aβ by APOE is dramatically mitigated [10, 11]. Hence, APOE isoform and its lipidation status determine its efficacy to eliminate Aβ in the brain.

Alu elements, primate-specific elements in humans, constitute over 10% of human genome and belong to a family of retroelements, termed short interspersed elements (SINEs). In contrast with human, mouse harbors two major SINEs, that is, B1 and B2 RNAs. B1 RNAs, similar as Alu RNA, deriving from 7SL RNA precursor, do not repress transcription whereas B2 SINEs, deriving from transfer RNA, repress polymerase II-dependent transcription[12, 13]. A plethoral of data indicate that Alu elements regulate gene expression by impacting polyadenylation[14], splicing [15], and affecting ADAR (adenosine deaminase acting on RNA) editing[16] or serving as transcriptional promoter or enhancer[17, 18]. Notably, Alu and B2 RNAs block or decrease transcription via disrupting the interaction between RNA polymerase II and target promoter[13, 19] or forming secondary structure to slow down migration of RNA polymerase II along mRNA[20]. When present in 3’-UTR, Alu RNA can form imperfect base-pairing with cytosolic Alu RNA and long non-coding RNA to transactivate Staufen1-mediated mRNA decay[21].

Dicer1 is a multidomain RNaseIII-containing enzyme to recognize and cleave 50-70-nucleotide hairpin structured pre-miRNAs or double stranded RNAs (dsRNAs) into functional miRNA or siRNA, respectively. In addition, Alu RNAs are also degraded by Dicer1[22] which is widely expressed in neuron, glia, and retinal pigment epithelial cells (RPEs); depletion of Dicer1 underlies neurodegeneration and degeneration of RPEs[22, 23]. Our previous study indicates that Dicer1 is reduced in the brain of an animal model of AD mice APPswe/PSEN1deltaE9 which is via Aβ42-mediated repression of NF-E2-related factor-2[24]. In this study, we indicate that delivery of Dicer1-expressing adenovirus improved spatial learning in the AD mice which was associated with enhanced clearance of amyloid peptide. Further, we found that Dicer1 increased ApoE expression by blocking inhibitory effect of Alu/B2 SINEs on ApoE.

## 2. Materials and methods

### 2.1. Materials

The following primary antibodies were used in this study:Dicer1(Sigma, St Louis, MO, USA, cat# WH0023405M1, RRID:AB_1841286), GFAP(Proteintech, Wuhan, China, cat#16825-1-AP, RRID:2109646), ApoE(Thermo Fisher Scientific, Carlsbad, CA, USA, Cat#701241, RRID:AB_2532438), Tubulin(Proteintech, Cat#14555-1-AP, RRID:AB_2212258), Aβ(1-16)(BioLegend, San Diego, CA, USA, Cat# 803001, RRID: AB_2721329). The following secondary antibodies were also used in this study:goat anti-mouse horseradish peroxidase conjugated IgG (Boster Biological Technology, Co. Ltd, Wuhan, China), goat anti-Rabbit horseradish peroxidase conjugated IgG(Boster Biological Technology). The following reagents or cell lines were used in this study:the Dicer1 siRNA duplex and the negative control(NC) siRNA duplex (Genepharma, Suzhou, China), pfu High fidelity enzyme(Qiagen, Beijing, China, cat#KP202), pJet1.2 vector(Thermo Fisher Scientific, Carlsbad, CA, USA, cat# K1231), pGL6-miR-basic (Beyotime Biotechnology, Haimen, China), pRL-TK Vector(Beyotime Biotechnology, Inc, cat#D2762), a human pCMV-Dicer 1(Sino Biological, Beijing, China, cat# HG11350-NH), U87MG cell line(ATCC, Manassas, VA cat#HTB-14, RRID:CVCL_0022).

### 2.2. Animal

All mice were fed water and food *ad libitum* in a standard pathogen-free facility with an automatic illumination on a 12-h on/off cycle in Wenzhou Medical University. All experiments were conducted complying with the ARRIVE guidelines and were carried out in accordance with the U.K. Animals(Scientific Procedures) Act, 1986 and associated guidelines, EU Directive 2010/63/EU for animal experiments and the protocols used were approved by the Wenzhou Medical University Committee on the Use and Care of Animals(wydw2020-0056). APPswe/PSEN1dE9(APP/PS1) mice(Jackson Laboratory, stock number 004462) were multiplied and genotyped according to the guidance by Jackson Laboratory. Both genders of transgenic mice were used and WT littermates were used as control mice. Behavioral experiments were performed during the daytime and at the same time of each day.

### 2.3. Intrahippocampal injection of Dicer1-expressing adenovirus

The adenovirus Ad-pCMV-EGFP(vehicle virus) or Ad-pCMV-Dicer1:T2A:EGFP virus expressing Dicer1 were packaged and generated by Cyagen Biosciences Inc (Guangzhou, China). Intraperitoneal injection of ketamine/xylazine(0.1/0.05 mg/g body weight) was used to anaesthetize mice which were then mounted on a stereotactic frame(KOPF, KD Scientific) for injection. With injection coordinates:anteriorposterior,-2 mm, mediolateral,± 2mm, dorsoventral,-2mm referring to bregma, the virus were injected into bilateral hippocampus with 1.5μl of viral titre(1.2 × 10^9^ vg/mL) in each hemisphere using a 10-μl Hamilton syringe(Hamilton Medical, Reno, NV, USA) connected to a 30-gauge micropipette, at an injection speed of 0.2 μl/min. The injections were conducted in 10.5-/11-month-old mice and behavior test was performed during 19-24 days after injection in which learning curve was acquired during 19-24 days and probe trial test conducted at the 25th day after injection. A quarter of mice were used for immunohistochemistry, quarter of them used for western blot, Aβ ELISA or quantitative real-time PCR, respectively.

### 2.4. Behavior test with Morris water maze

Spatial memory was tested by use of a Morris water maze and three groups of mice, WT mice(with intrahippocampal injection of Ad-pCMV-EGFP virus) and APP/PS1 mice (intrahippocampal injection of Ad-pCMV-EGFP or Ad-pCMV-Dicer1-T2A:EGFP) were tested. A nontoxic white paint was added to water (24°C) in the water maze which was surrounded by a black curtain(placed 80 cm away) that held three salient visual cues. Initially, mice were randomly trained in four quadrants of the maze, and trained to find a hidden platform(10-cm diameter, 1 cm under the water surface) with a maximum of 60 s. Before probe trial test, mice were trained for 4 trials per day, 10-min interval between trials for 6 days. On day 7, mice were given a 60-s probe test to find the platform. Swimming pattern, distance, speed, and the amount of time spent in each of the four quadrants were recorded with a SLY-WMS Morris Water Maze System (Beijing Sunny Instruments Co. Ltd) consisting of a frame grabber, a video camera, a water pool, and an analysis software.

### 2.5. Western blot analysis

The mice used were fasted overnight, and at the next morning, mice were anesthetized by isoflurane and transcardially perfused with ice-cold phosphate buffer (PBS). The hippocampi were isolated and freshly frozen in liquid nitrogen. For analyzing the U87MG cells transfected by pCMV-Dicer1 plasmid, the cells were plated at the density of 4×10^6^ cells/well onto six-well plates. For analyzing the cells infected by virus expressing Dicer, primary cultured astrocytes were seeded in six-well plates at the density of 5X10^5^ cells / well. The examined tissues or cell samples were homogenized in RIPA buffer composed of components in mM:460 Tris–HCl, pH 7.4, 138 NaCl, 1 EDTA, 2.5 NaF, 2.5 Na3VO4, 1 phenylmethanesulfonylfluoride, 1 dithiothreitol, supplemented with 0.1% Nonidet P-40, and 1Xprotease/1Xphosphatase inhibitor cocktail(Sigma, St.Louis, MO). Proteins were resolved on 12% SDS-PAGE gels(Bio-Rad) and transferred to nitrocellulose membranes(ThermoFisher). Membranes were blocked in blocking buffer(Beyotime Biotechnology) and then incubated with primary antibody. Following wash, the membranes were then incubated with secondary antibody, and developed with chemiluminescence system kit(ThermoFisher). The blots were quantified using ImageJ software(National Institutes of Health, Bethesda, MD, USA).

### 2.6. Quantitative real-time polymerase chain reaction (qRT-PCR)

The hippocampi were isolated and freshly frozen in liquid nitrogen. U87MG cells were plated on 6-well plate until ~70% confluency, and transfected with 1.5 μg pCMV3 Dicer 1 vector or pCMV empty vector, or transfected with human Dicer1 siRNA or AluSz RNA for 48 h in sera-free DMEM. The cultured primary mouse astrocytes were plated onto 6-well plate until ~50% confluency and cells were infected with adenovirus or transfected with mouse Dicer 1 siRNA for 60 h before analyses. Extraction of total RNAs from hippocampi or transfected cells and elimination of genomic DNA were performed by a RNA extraction kit(TIGEN BIOTECH, Co, LTD., Beijing, China, cat# DP424). RNA (200 ng) was used to synthesize the cDNA by a first stand cDNA synthesis kit(cat#R111-02, Vazyme, NanJing, China). Fifty nanograms of cDNA were used as a template to amplify individual mRNA. The following primers were used in qRT-PCR:*hDicer 1*, sense, 5’-CCCGGCTGAGAGAACTTACG-3’, antisense, 5’-CTGTAACTTCGACCAACACCTTTAAA-3’;*hApoE*, sense, 5’-GTTGCTGGTCACATTCCTGG-3’, antisense, 5’-GCAGGTAATCCCAAAAGCGAC-3’; AluSz downstream of the 3’-UTR of human ApoE mRNA,*s*ense, 5’-CAACATAGTGAAACCCCGTCTCT-3’; antisense, 5’-GCCTCAGCCTCCCGAGTAG-3’; The 18srRNA, sense, 5’-TTCGTATTGCGCCGCTAGA-3’, antisense, 5’-CTTTCGCTCTGGTCCGTCTT-3’;*mDicer1*, sense, 5’-GTCAGCCGTCAGAACTCACTC-3’, anti-sense, 5’-ACAGTCAAGGCGACATAGCAA-3’;*mApoE*, sense, 5’-GACCCAGCAAATACGCCTG-3’; antisense, 5’-CATGTCTTCCACTATTGGCTCG-3’; *B2Mm2* RNA, sense, 5’-GGTGCTGGAGAGATGGCTCA-3’, anti-sense, 5’-AGATTTGTTTATCTTATGT-3’. The SYBR™ Green(Invitrogen) was used as the dye for the qRT-PCR reaction following the manufacturer’s instruction. The amplification processes were initiated at 50 °C for 2 min and an ensued step at 95 °C for 10 min, ended with 40 cycles of PCR reactions(95 °C for 10 s and 60 °C for 1 min). Reaction specificity, indicated by the presence of a single amplification peak for each PCR reaction, was examined by use of dissociative curves. 2^-ΔΔCt^ method was used to quantify the reaction, which was normalized to corresponding control groups.

### 2.7. ELISA assay for soluble/insoluble Aβ

The soluble form of human Aβ40 and Aβ42 contained in the hippocampus from APP/PS1 mice were extracted with diethylamine(DEA, CAS No.109-89-7, Cat# D110465, Aladdin Co., Ltd, Shanghai, China) and the insoluble form were extracted with formic acid (FA, CAS No.64-18-6, Cat#F112031, Aladdin Co., Ltd) according to a described protocol (http://www.bio-protocol.org/e1787). Human Aβ40 and Aβ42 were individually measured in DEA (soluble Aβ) and FA (insoluble Aβ) fractions using commercially available kits (Cat#CSB-E08299h for human Aβ40; Cat#CSB-E10684h for human Aβ42, CUSABIO TECHNOLOGY LLC, Wuhan, China) per the manufacturer’s instructions. These kits utilize neoepitope-specific antibodies(Abs) to specifically detect either human Aβ40 or Aβ42 and have no cross-reactivity with full-length APP.

### 2.8. Immunofluorescence

The 8- or 11-month-old WT orAPP/PS1 mice were deeply anesthetized by isoflurane, killed, and then transcardially perfused with ice-cold PBS followed by perfusing with 4% paraformaldehyde (PFA) in PBS. The isolated brains were postfixed in 4% PFA overnight, stored in PBS containing 0.01% sodium azide. The brains were mounted in agarose and 20 μm coronal sections were obtained using a Leica VT1000S Vibratome. Antigen retrieval was performed in 0.01M sodium citrate solution (pH 6.0) at 99°C for 40 min. After washed in PBS three times, 5 min each, at room temperature (RT), the sections were blocked using 5% normal donkey serum in PBS containing 0.2% Triton X-100 for 1 h at room temperature. The sections were individually incubated with monoclonal anti-Dicer1(1:300, Sigma-Aldrich), mouse anti-GFAP(1:1000, Proteintech), mouse anti-Aβ(1-16) antibody(1:800, Biolegend), or mouse ant-ApoE antibody(1:800, Cell signaling)overnight at 4°C. For mice injected with virus, the sections were incubated with anti-Aβ(1:300, Sigma-Aldrich) or anti-GFAP(1:1000, Proteintech). After washed 3 times with PBS (3×5 min at RT), the sections were incubated with secondary antibodies: Alexa Fluor 594 goat anti-rabbit or Alexa Fluor 488 goat anti-mouse IgG(1:1000, Life Technologies Corporation, Carlsbad, CA, USA) for 1 h at RT, then washed three times with PBS (3×5 min at RT). The sections were then mounted on coverslips(Citotest Labware Manufacturing Co., Ltd, Jiangsu, China) and the images of hippocampus were obtained with a fluorescent microscope(DMi8, Leica Biosystems).

### 2.9. Fluorescence *in situ* hybridization

Mouse primary cultured astrocytes were plated at the density of 1.5X10^5^ cell per dish in 3.5 cm Falcon^®^ dishs(Cat#353001, BD Biosciences). After transfected with mouse Dicer1 siRNA for 12 h, the cells were fixed by 4% PFA for 10 min at RT. The cells in dish were incubated with 0.1% TritonX-100 in PBS containing 2% diethyl pyrocarbonate for 30 min at RT, and washed with 2Xsaline sodium citrate(SSC) buffer. Cy3-probe were mixed with hybridization buffer and added to the cultured cells and incubated overnight at 37°C according to the manufacturer’s instruction(Cat#F05401, GenePharma Co., Ltd, Soochow, China). After washing, mouse anti-Dicer1 antibody(1:500 in PBS) was added into dish and incubated with the cells for 8 h at 4 °C. After wash, the cells were incubated with secondary antibody, Alexa Fluor488 goat anti-mouse IgG (1:1000), at 37°C for 1 h. The 2-(4-Amidinophenyl)-6-indolecarbamidine dihydrochloride (DAPI, Beyotime) diluted in PBS at 100 μg/ml was used to stain nuclei at RT for 5 min. The signals of Dicer 1 and B2 Mm2 RNA and images of cells were obtained with a fluorescent microscope(DMi8, Leica Biosystems). The following probes were used:B2 Mm2 RNA probe 2, 5’-AACCACATGGTGGCTCACAACCAT-3’; negative control probe, 5’-TGCTTTGCACGGTAACGCCTGTTT-3’; positive 18s RNA probe, 5’-CTGCCTTCCTTGGATGTGGTAGCCGTTTC-3’.

### 2.10. Northern blot

The hippocampi of WT or APP/PS1 mice at age of 11 months were freshly frozen in liquid nitrogen, respectively. The cultured primary mouse astrocytes were plated into 6-well plate until ~50% confluency and transfected with mouse Dicer 1 siRNA for 48 h. Extraction of total RNAs from hippocampi or transfected cells was performed by a kit (TIGEN BIOTECH, Co, LTD., cat# DP424, China). The isolated RNAs were loaded, separated and transferred onto nitrocellulose filter membrane(Beyotime, cat#FFN02, China) following by UV crosslink. All procedures were performed according to the instruction from Northern™ Max kit(ThermoFisher, cat#AM1940). Following wash, the membrane was incubated with 3’-biotinylaion probes(Stargene Co., Ltd, Wuhan, China). The following synthesis 3’-end biotin-probes were used:B2 Mm2 RNA probe, 5’-AGCACTGACTACTCTTCTATAAGTC-3’. Mouse U6RNA probe, 5’-CAGCACAAAAGGAAACTCACC-3’. The membrane was incubated with secondary streptavidin-HRP antibody(Beyotime, 1:800, cat# A0303). The membranes were developed with chemiluminescence system kit(Thermo Fisher).

### 2.11. Astrocyte cell cultures

The brains from mice at day 1 after birth were used for primary astrocyte cell culture. The brains were isolated and digested in 2 ml trypsin (0.25%) for 15 min at 37°C, which was then inactivated by addition of 3 ml of 10% fetal bovine serum (FBS). The tissue was triturated by a pipette to make a homogenous mixture, which was passed through a cell strainer to remove un-dissociated tissue. The cells were centrifugated for 5 min at 1, 800× *g*, and the supernatant was discarded. The cell pellets were resuspended in complete DMEM media containing antibiotics(1% penicillinstreptomycin mixture) and 20% FBS. The cells were then plated in a culture plate and continued culturing for 20 days. Before transfection, the medium was replaced with DMEM/F12 containing 0.1% bovine serum albumin. After 24 h in culture, astrocyte cultures were used for transfection or virus infection. U87MG cells, a human astrocytoma cell line, were cultured in Dulbecco’s modified Eagle’s Medium(DMEM) containing 10% FBS.

### 2.12. Plasmid construction and Luciferase reporter assay

With genomic DNA extracted from human U87MG cells as a template, the human ApoE 3’-untraslated region (3’-UTR) and its 3’-flanking inverted Alu-like sequence: AluSz and the following two AluJo(Entrez acc.no.:NM_001302688.2) were amplified to produce an amplicon which spanned ApoE mRNA 3’-UTR starting from stop code (0 bp) and its downstream 1160bp, termed 3’ UTR-AluSz-2XAluJo. The PCR reaction was conducted with forward primer 5’-CGACGCGTTGCCCAGCGACAATCAC-3’ and reverse primer 5’-CCCAAGCTTAGGAGGGAGAGATAAAAGA-3’. The construct with deletion of the 2^nd^ AluJo, 3’-UTR-AluSz-AluJo, was amplified by alternating reverse primer 5’-CCCAAGCTTGTGTGGTGGTGCTAGCCA-3’ to produce a 874-bp PCR product whereas the 3’-UTR-AluSz with deletion of two AluJo was amplified by alternating reverse primer 5’-CCCAAGCTTAAGAAAGAGGGGCA-3’ to produce a 583-bp PCR product. The construct not containing inverted Alu SINEs was also amplified by alternating reverse primer 5’-CCCAAGCTTGGGCAGAGAGAAAGATA-3’, producing a 210-bp PCR product (3’-UTR). For generating construct containing mouse 3’-UTR and its 3’-flanking B2 Mm2 RNA derived from mouse ApoE mRNA (Entrez acc.no.: NM_00009696.4), the genomic DNA was extracted from primary mouse astrocyte cultures. The PCR reaction was conducted using forward primer 5’-CGACGCGTGTATCCTTCTCCTGTCCT-3’ and reverse primer 5’-CCCAAGCTTTAGAAGAGCCTGTACTGGGTG-3’, producing an amplicon spanning mouse ApoE mRNA 3’-UTR and its downstream 679 bp sequence, termed 3’-UTR-B2Mm2. The 3’-UTR from mouse ApoE was amplified by alternating reverse primer 5’-CCCAAGCTTTGGCTCAGTGGTAGATCAC-3’. All PCR products were amplified by pfu High fidelity enzyme and cloned into a pJet1.2 vector(ThermoFisher) and then subcloned into a luciferase reporter, pGL6-miR-basic(cat# D2106, Beyotime, Haimen, China). The constructs spanning human ApoE 3’-UTR or/ and its flanking Alu/B2 SINEs were referred as pGL6-luc-3’-UTR, pGL6-luc-3’-UTR-AluSz, pGL6-luc-3’-UTR-AluSz-AluJo, and pGL6-luc-3’-UTR-AluSz-2XAluJo. The constructs spanning mouse ApoE 3’-UTR or/and its flanking inverted B2 SINE were referred as pGL6-3’-UTR and pGL6-luc-3’-UTR-B2Mm2. For convenient subcloning, all forward primers incorporating Mlu I restriction site and reverse primer containing HindIII restriction site were used to generate product and subcloned into PGL6-miR-basic.

The U87MG cells were seeded at the density of 3X10^4^ cells/per well in 48-well plate and transfected according to the manufacturer’s protocol(Lipofectamine 2000, Invitrogen). The *renilla* luciferase-containing reporter(pRL-TK Vector) was mixed with individual construct containing *firefly* luciferase, and a human Dicer1 siRNA or an scrambled negative control siRNA at a ratio of 1:20:100 in mass, or co-transfected with synthesized AluSz RNA at a ratio of 1:20:10 in mass, respectively. For mouse ApoE 3’-UTR luciferase activity assay, the primary cultured astrocytes were plated into 48-well plate at 2X10^4^ cells/ per well. The *renilla* luciferase-containing reporter(pRL-TK Vector) was mixed with individual construct containing *firefly* luciferase, and a mouse Dicer siRNA or a scrambled negative control siRNA at a ratio of 1:20:100 in mass, or cotransfected with synthesized B2 Mm2RNA at a ratio of 1:20:20 in mass, respectively. The luciferase assay was conducted in a plate reader(SpectraMax M5, Molecular devices, San Jose, CA, USA) per the manufacturer’s protocol(Beyotime Biotechnology, cat#RG027). The values were recorded as relative luminescence unit(RLU) between *firefly /renilla* luciferase.

### 2.13. Syntheses of ApoE AluSz/B2 Mm2 RNA

For synthesis of ApoE AluSz RNA, the genomic DNA was extracted from U87MG cells and amplified with PCR reaction with forward primer:5’-<u>TAATACGACTCACTATAG</u>GGCCCCGTCT-3’ containing underlined T7 RNA polymerase promoter and reverse primer:5’-GAGACGGGAAAAAAAAAAAAT-3’. With PCR production as a template, AluSz RNA was synthesized under the guidance of the T7 RNA promoter in a reaction buffer containing T7 RNA polymerase(cat#C11002-1 RiboBio Co., Ltd, Guangzhou, China), NTP, RNase inhibitor, and DEPC-treated water. The product was extracted by gel purification and produced transcripts, yielding single strand RNAs that fold into a defined secondary structure, identical to RNA polymerase III-derived transcripts. For synthesis of AluSz RNA antisense, the genomic DNA was amplified with forward primer:5’-<u>TAATACGACTCACTATAG</u>GGCTCTGCCCTT-3’ containing underlined T7 RNA polymerase promoter and reverse primer:5’-GGGGCAGAGGCCGGGCAT-3’. With the PCR product as a template, the antisense RNA was also synthesized with T7 RNA polymerase. For synthesis of mouse ApoE B2 Mm2 RNA, genomic DNA was extracted from mouse brain and amplified with forward primer:5’-<u>TAATACGACTCACTATAG</u>GGGGTGCTGGAGAGAT-3’ and reverse primer: 5’-GGATTCTAGGCATGGGCTCATT-3’. Similarly, the template for B2 Mm2 RNA antisense RNA was amplified with forward primer:5’-TAATACGACTCACTATAGGGGGATTCTAGGCATG-3’ and reverse primer:5’-<u>TAGAAGAGCCTGTACTG</u>GGTGCT-3’. All the transcription reaction was conducted with T7 RNA polymerases according to the manufacturer’s recommendation(RiboBio Co., Ltd, Guangzhou, China). The synthesized *Alu/B2* RNA were precipitated, suspended in water and analyzed on 1.5% non-denaturing agarose gel, respectively. All antisense RNAs were set as negative control in following transfection experiments.

### 2.14. Experimental design and statistical analysis

During Water Maze test, mice were evenly allocated to each experimental group according to the gender and ages, and the tests were conducted in a blinded manner, that is, the investigator was blinded to the group allocation of the genotype or treatment during the experiment and when assessing all the results. All the data acquired during experiments were included for analysis and there were no sample size differences between the beginning and end of experiments. All data were presented as means ± SEM unless specified. Single data points were shown as overlaying dot-plot on bargraph when sample sizes were smaller than 15, otherwise, shown as bar-graph. For parametric data, Student’s *t*-test was used for comparing differences between two groups. In experiments with more than two groups, one-way ANOVA was performed followed by Tukey’s *post hoc* test for comparisons among groups. For analysis of two groups of non-parametric data, the Mann-Whitney *U*-test was used for determining the differences. For Morris water maze experiment, two-way ANOVA with repeat measures followed by Tukey’s *post hoc* test was used for analyzing the time spent in four quadrants, frequency to crossing platform, swimming speed and distance. Statistical analyses were performed with Graphpad Prism 7.04 software(Graphpad software, Inc. La Jolla, CA, USA). Difference was regarded as significance when *p*<0.05.

## 3. Results

### 3.1. Overexpression of Dicer1 improved spatial learning in APP/PS1 mice

Our previous study indicates that injection of Dicer1-expressing virus in the hippocampus of APP/PS1 mice at the age of ~4 months improves learning, relative to injection of vehicle virus[24]. Herein, we aimed to examine the beneficial effect in ~11-month-old APP/PS1 mice which develop widespread deposition of amyloid plaques as reported [25]. During the training period on day 6 after virus injection, the time to reach the platform was longer for APP/PS1 mice than WT mice (p=0.0162); the effect was reversed by injection of Ad-pCMV-Dicer1:T2A:EGFP compared with injection of Ad-pCMV-EGFP on day 6(p=0.0148). In probe trial test on day 7, the animals’ swimming pattern, distance, speed, and the time spent in each quadrant of swimming pool were recorded. Without differences on swimming distance (Figure 1D) and swimming speed (Figure 1E), APP/PS1 mice spent less time in target quadrant than WT mice in probe trial test(p=0.02) (Figure 1B), which was reversed by injection of Ad-pCMV-Dicer1:T2A:EGFP compared with injection of Ad-pCMV-EGFP(p=0.0423) (Figure 1B). Similarly, APP/PS1 mice crossed the hidden platform less frequently than WT mice (p=0.0117), which was increased by injection of Ad-pCMV-Dicer1:T2A:EGFP virus, relative to the effect by injection of Ad-pCMV-EGFP virus.

**Figure 1.**
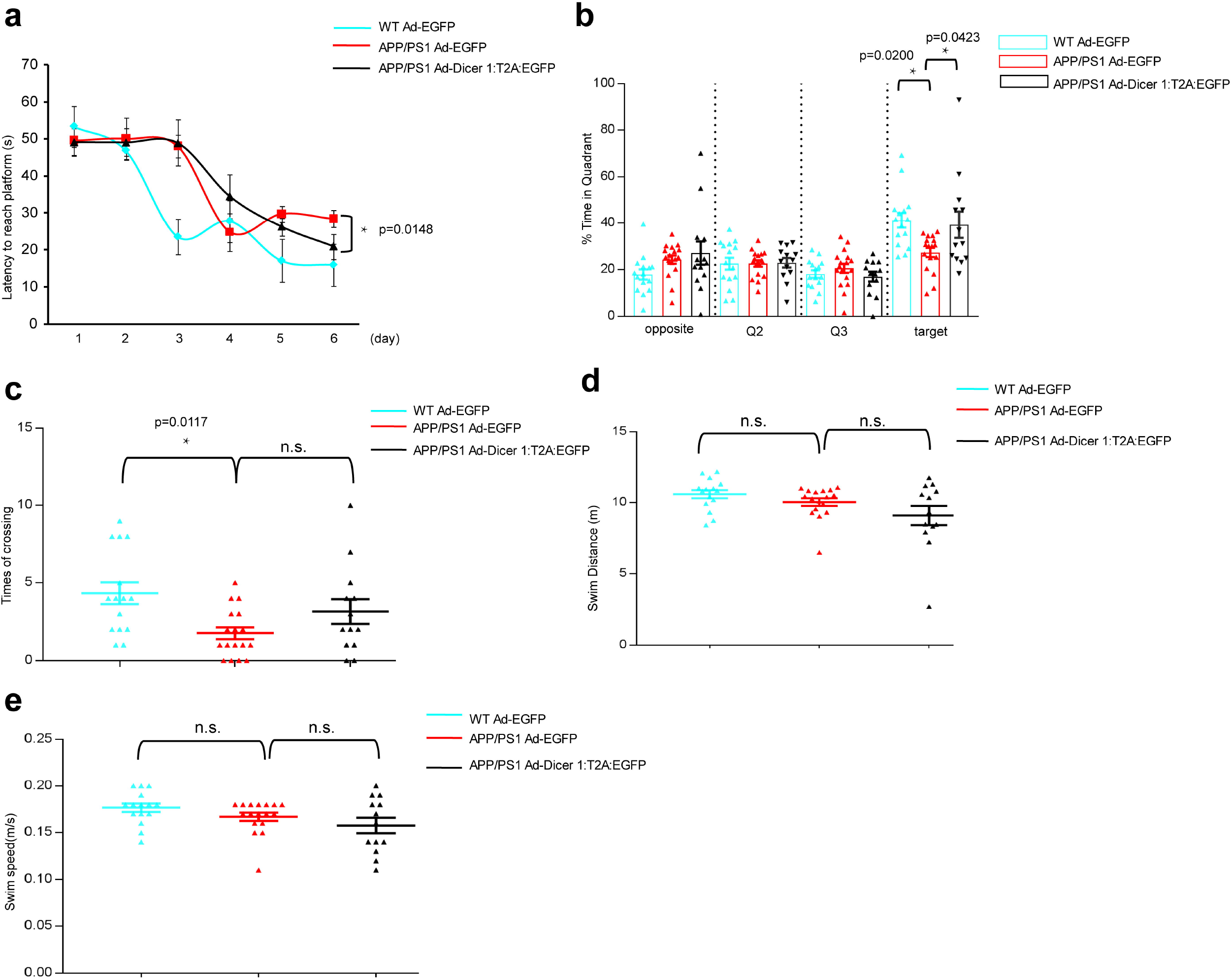
Intrahippocampal injection of Dicer1-expressing adenovirus partially but significantly improved spatial learning in ~11-month old APP/PS1 mice. Ad-pCMV-Dicer1-T2A:EGFP virus (1.2 × 10^9^ vg/mL) or equal amount of vehicle virus Ad-EGFP virus were injected into CA3 hippocampus of ~11-month-old APP/PS1or WT mice and behavior test was conducted at 19-24 days after injection. **(a)** Learning curve from WT mice injected with Ad-pCMV-EGFP virus (WT-Ad-EGFP, n=15), APP/PS1 mice injected with Ad-pCMV-EGFP virus (APP/PS1 Ad-EGFP, n=17) or Ad-pCMV-Dicer1-T2A:EGFP virus (APP/PS1 Ad-Dicer1:T2A:EGFP, n=13). The latency to reach platform were analyzed by use of two-way ANOVA with repeated measures against day and treatment (virus injection), F_interaction_ (10, 225) =1.62, p=0.1011; F_treatment_ (2, 225) =6.695, p=0.0015; F_day_(5, 255)=24.16, p<0.0001. Tukey’s *post hoc* test was further used to compare the differences between APP/PS1 mice injected with Ad-pCMV-Dicer1:T2A:EGFP virus and injected with Ad-pCMV-EGFP virus, * p=0.0148 on day 6. **(b)** Preference of APP/PS1mice for the target quadrant was enhanced by injection of Ad-pCMV-Dicer1:T2A:EGFP virus in CA3 hippocampi. Two-way ANOVA with repeated measures against quadrant and treatment was used to compare the differences, F_interaction_(6, 126)=2.879, p=0.0116; F_treatment_(2, 42)=2.725, p=0.0772, F_Quadrant_ (3, 126)=18.89, p<0.0001. Tukey’s *post hoc* test was further used to compare the differences among groups, APP/PS1 Ad-Dicer1:T2A:EGFP (n=13) *vs*. APP/PS1 Ad-EGFP(n=17), * p=0.0423; APP/PS1 Ad-EGFP (n=17) *vs*. WT Ad-EGFP (n=15), * p=0.02. **(c)** Frequencies of crossing target platform was increased by injection of Dicer1-expressing virus. One-way ANOVA was used to compare the differences and Tukey’s *post hoc* test was further used to compare the differences among groups, F(2, 42)=4.588, p=0.0158, APP/PS1 Ad-Dicer 1-T2A:EGFP (n=14) *vs*. APP/PS1 Ad-EGFP (n=15), n.s; APP/PS1 Ad-EGFP (n=15) *vs*. WT Ad-EGFP (n=16), * p=0.0117. **(d)** Swim distance and **(e)** Swim speed did not show differences among groups (n.s).

### 3.2. Overexpression of Dicer1 reduced β-amyloid deposition, increased APOE, and reduced B2 Mm2 RNA in APP/PS1 mice

To investigate the mechanism by which expression of Dicer1 in the hippocampus improved spatial learning, we examined β-amyloid in hippocampus by immunofluorescence and ELISA examination. With immunofluorescence, we found that APP/PS1 hippocampus contained more stainings against amyloid, mostly in stratum radiatum, than WT mice; APP/PS1 mice injected with Ad1-pCMV-Dicer1:T2A:EGFP contained lower numbers of and smaller-size stainings than did APP/PS1 mice injected with vehicle virus, Ad1-pCMV-EGFP (Figure 2A). As a positive control, injection of Ad1-pCMV-Dicer1:T2A:EGFP in hippocampus did not decrease β-amyloid stainings in the parietal lobe in APP/PS1 mice compared with the AD mice injection with vehicle virus in hippocampus (Figure 2B). However, the parietal lobe from APP/PS1 mice contained much higher numbers of amyloid stainings compared with WT mice (Figure 2B). We further examined β-amyloid content by ELISA with hippocampal homogenates. Soluble and insoluble Aβ42 from the homogenates of APP/PS1 mouse hippocampus were higher than that from WT mice, respectively; both forms of Aβ42 were significantly reduced by injection of Ad1-pCMV-Dicer1:T2A:EGFP compared with injection of vehicle virus (p=0.0002 for soluble, and p=0.0006 for insoluble Aβ42), respectively (Figure 2C). Similarly, insoluble Aβ40 from the homogenates of APP/PS1 mouse hippocampus was higher than that from WT mice which was significantly reduced by injection of Ad1-pCMV-Dicer1:T2A:EGFP compared with injection of vehicle virus(p=0.0022) (Figure 2C). By contrast, the contents of soluble Aβ40 were not different among the hippocampus of the three groups of mice (Figure 2C). We continued to conduct western blot against Dicer1, APOE, and β-amyloid with hippocampal homogenates. As expected, injection of Dicer1-expressing adenovirus increased Dicer1 protein in APP/PS1 hippocampus which contained less Dicer1 than WT mice (Figures. 2D and 2E), consistent with our previous report that Dicer1 was reduced in APP/PS1 hippocampus relative to WT[24]. APOE protein levels in the hippocampus of AD mice were not different with WT mice; expression of Dicer1 though increased APOE level in APP/PS1 hippocampus compared with injection of vehicle virus (Figures.2D and 2F, p=0.0002). We also tested APOE protein levels in the hippocampus of APP/PS1 mice or WT mice at the age of 4 months upon injection of Dicer1-expressing virus or vehicle virus(Supplemental Figure 1 and 2). Similarly, injection of Dicer1-expressing virus into hippocampus increased APOE protein level in AD hippocampus compared with injection of vehicle virus; the results were examined with western blot(Supplemental Figure 1) and immunofluorescence assay (Supplemental Figure 2). In accordance with immunofluorescence and ELISA analysis of β-amyloid, APP/PS1 hippocampus contained higher amount of β-amyloid than WT (Figures. 2D and 2G, p<0.0001) which was reduced by injection of Dicer1-expressing virus compared with injection of vehicle virus (Figures. 2D and 2G, p<0.0001). Notably, astrogliosis were found in AD mice compared to WT mice(Supplemental Figure 3) and injection of Ad1-pCMV-Dicer1:T2A:EGFP reduced astrogliosis in the hippocampus of AD mice compared with injection of vehicle virus, Ad-pCMV-EGFP(Supplemental Figure 4). Since B2 RNA is a substrate of Dicer1, we examined the content of B2 Mm2 RNA in APP/PS1 mice. With northern blot, B2 Mm2 RNA was increased in the hippocampus of AD mice relative to that from WT mice (Figures. 3A and 3B, p=0.0022). Consistent with our previous result[24], Dicer1 mRNA was reduced in the hippocampus of ~11-month-old APP/PS1 mice compared with WT mice at similar age(p=0.0418); injection of Dicer1-expressing virus increased Dicer1 mRNA in the hippocampus of APP/PS1 mice(p=0.0475) (Figure 3C). In line with the reduction of Dicer1 in AD hippocampus, B2 Mm2 RNA was increased in the hippocampus of AD mice related with that of WT mice (p=0.0069); injection of Dicer1-expressing virus decreased B2 Mm2 RNA in the hippocampus of APP/PS1 mice(p<0.0001) (Figure 3D). Interestingly, it was found that *APOE* mRNA was not different between the hippocampus of the AD mice and that of WT mice(n.s); however,*APOE* mRNA was increased in the hippocampus of the AD mice by injection of Ad1-pCMV-Dicer1:T2A:EGFP versus injection of vehicle virus(p=0.0027) (Figure 3E).

**Figure 2.**
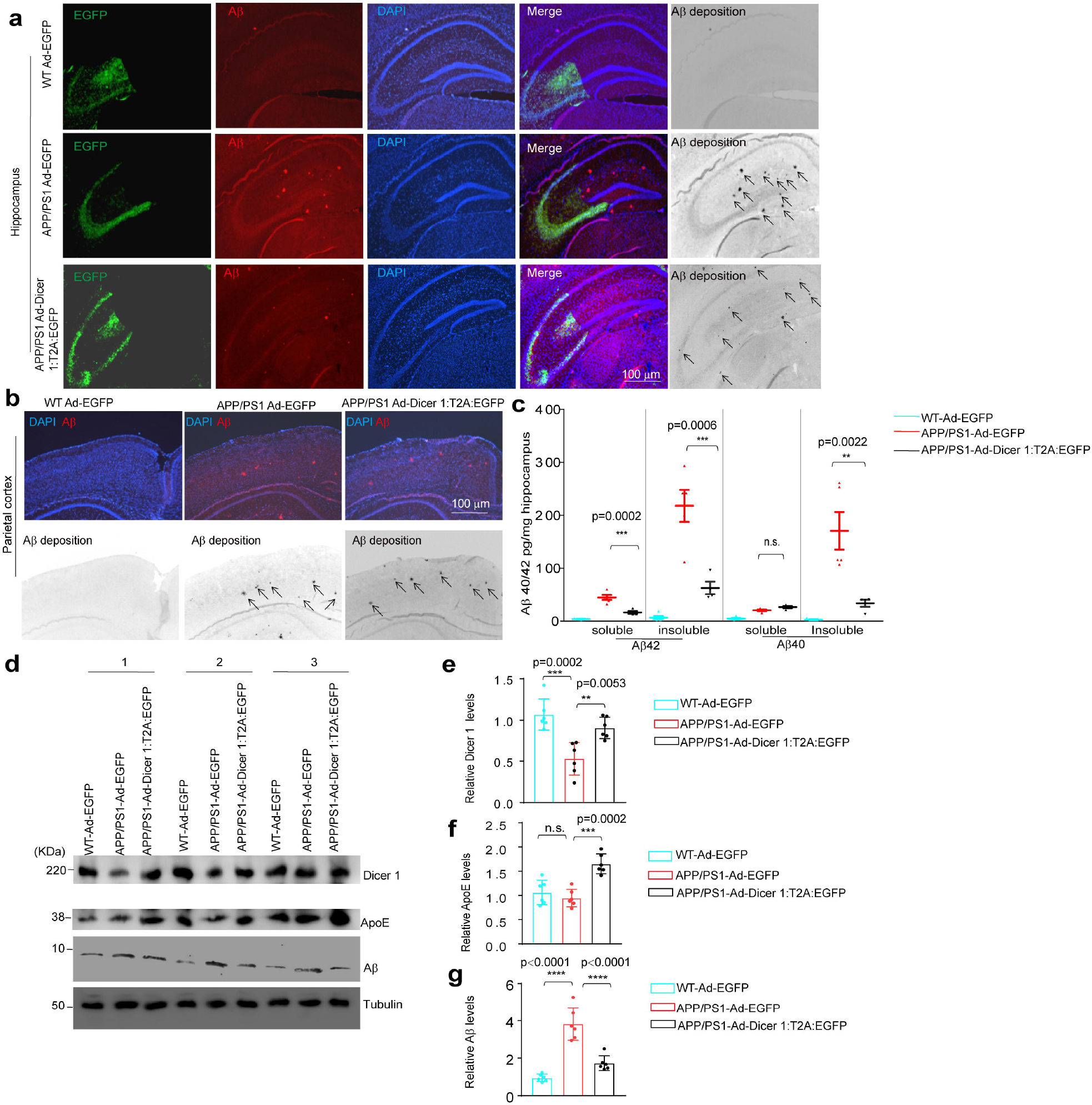
Dicer 1 overexpression enhanced APOE expression and promoted Aβ clearance in the hippocampus of APP/PS1 mice. WT or APP/PS1 mice at ~11 months were injected with 2 μL of Ad-pCMV-EGFP or Ad-pCMV-Dicer1:T2A:EGFP virus (1.2 × 10^9^ vg/mL) in hippocampus and parietal lobe as described in the materials and methods. After 25 days, the mice were subject to decapitation and the brain was processed into 10 μm sections, fixed in 4% paraformaldehyde and stained for Aβ and DAPI. The images were captured with fluorescence microscope with different fluorophore filter for DAPI, GFP and red channel for cell nuclei, virus infection, and Aβ deposition, respectively. **(a)** The first column indicated virus infection; second column, Aβ stainings; the third column, cell nuclei; and the fourth column indicated the merged images of all the stainings. Aβ stainings were converted into phase-contrast images in the fifth column. Scale bar, 100 μm. **(b)** Signals of Aβ deposition in parietal cortex of WT mice and APP/PS mice injected with vehicle virus or Dicer1-expressing virus. Top row indicated the color stainings of Aβ (red) and DAPI (blue) and low row indicated Aβ staining in a phase-contrast mode. Scale bar, 100 μm. The arrows indicated Aβ deposition. **(c)** Injection of Dicer1-expressing virus reduced Aβ40 or Aβ42. Soluble/insoluble Aβ were extracted from hippocampus of virus-injected mice from each group. The concentrations of soluble or insoluble Aβ40, Aβ42 were measured by ELISA. One-way ANOVA was used to compare the differences and Tukey’s *post hoc* test was further used to compare the differences in mice injection with Dicer1-expressing virus (n=4) and vehicle virus (n=5), *** p=0.0002 for comparison of soluble Aβ42, *** p=0.0006 for comparison of insoluble Aβ42; n.s. for comparison of soluble Aβ40; ** p=0.0022 for comparison of insoluble Aβ40. The amount of Aβ40 or Aβ 42 in WT mice with virus injection were set as basal levels. **(d)** Injection of Dicer1-expressing virus increased APOE and reduced Aβ40, Aβ42 in the hippocampus of AD mice. The hippocampal tissues were extracted, homogenized, and subject to western blot examination of Dicer1, APOE, and Aβ. The loading protein was separated by SDS-PAGE and tubulin was used a loading control protein. The protein levels were normalized to WT Ad-EGFP groups accordingly. **(e)** Relative Dicer 1 protein levels: APP/PS1 Ad-Dicer1:T2A:EGFP (n=6) *vs*. APP/PS1 Ad-EGFP(n=6), ** p=0.0053; APP/PS1 Ad-EGFP (n=6) *vs*. WT Ad-EGFP (n=6), *** p=0.0002. **(f)** Relative APOE protein levels: APP/PS1 Ad-Dicer 1-T2A: EGFP (n=6) *vs*. APP/PS1 Ad-EGFP(n=6), *** p=0.0002; APP/PS1 Ad-EGFP (n=6) *vs*. WT Ad-EGFP (n=6), n.s. p= 0.6226. **(g)** Relative Aβ protein levels: APP/PS1 Ad-Dicer1:T2A:EGFP (n=13) *vs*. APP/PS1 Ad-EGFP(n=17), **** p<0.0001; APP/PS1 Ad-EGFP (n=17) *vs*. WT Ad-EGFP (n=15), **** p<0.0001.

**Figure 3.**
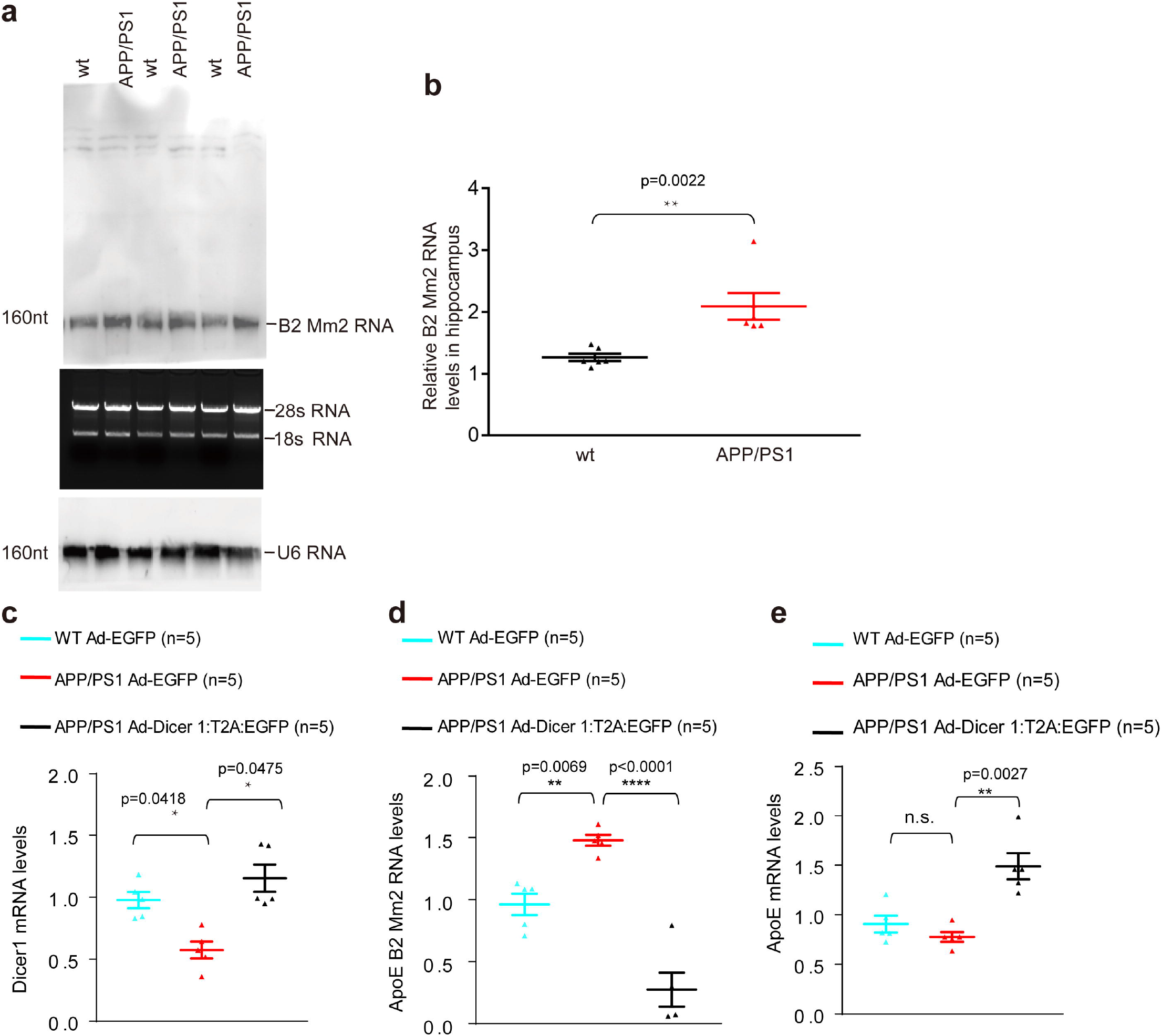
Dicer1 overexpression reduced B2 Mm2 RNA in the hippocampi of APP/PS1 mice. WT or APP/PS1 mice at ~11 months were injected with 2 μL of Ad-pCMV-EGFP or Ad-pCMV-Dicer1:T2A:EGFP virus (1.2 × 10^9^ vg/mL) in hippocampus as described in the materials and methods. After 19 days, the total RNA from hippocampus in each group was isolated to determine Dicer1 mRNA, *APOE* mRNA and B2 Mm2 levels. One-way ANOVA was used to compare the differences and Tukey’s *post hoc* test was further used to compare the differences among groups. **(a)** Injection of Ad-Dicer1:T2A: EGFP virus into the hippocampus of APP/PS1 (n=6) increased Dicer1 mRNA in the hippocampus compared with of APP/PS1 mice injected with Ad-EGFP virus (n=6), * p=0.0475; the hippocampus of AD mice injected with vehicle virus (n=6) contained less amount of Dicer1 mRNA than of WT mice injected with vehicle virus (n=6), * p=0.0418. The mRNA levels of Dicer1 were determined by qRT-PCR. **(b)** Injection of Ad-Dicer1-T2A:EGFP virus into the hippocampus of APP/PS1 (n=5) reduced B2 Mm2 RNA in the hippocampus compared with of APP/PS1 mice injected with Ad-EGFP virus (n=5), **** p<0.0001; the hippocampus of AD mice injected with vehicle virus (n=5) contained higher amount of B2 Mm2 RNA than of WT mice injected with vehicle virus (n=5), ** p=0.0069. The levels of B2 Mm2 RNA were determined by qRT-PCR. **(c)** Injection of Ad-Dicer1:T2A:EGFP virus into the hippocampus of APP/PS1 (n=5) increased *APOE* mRNA in the hippocampus compared with of APP/PS1 mice injected with Ad-EGFP virus (n=5), ** p=0.0027; the hippocampus of AD mice injected with vehicle virus (n=5) contained similar amount of *APOE* mRNA as WT mice injected with vehicle virus (n=5), n.s., p=0.5285. The *APOE* mRNA levels were determined by qRT-PCR. The primary cultured astrocytes were plated at the density of 1 × 10 ^5^ cells /per well in 6-well plate until ~50% confluency and transfected with Dicer1 siRNA and NC siRNA for 60 hours and the levels of Dicer 1 mRNA, *APOE* mRNA and B2 Mm2 RNA were determined by qRT-PCR. Paired Student’ *t-test* was utilized to compare differences between two groups.

### Reduction of Dicer1 induced accumulation of B2 Mm2 RNA and reduced APOE mRNA in astrocyte cultures

To investigate the mechanism, we examined the effect by knocking down Dicer1 on APOE in primary mouse astrocyte cultures. We have tested two sets of Dicer1 siRNA and chose the one with better efficacy; in U87MG cultures, Dicer1 knockdown reduced *APOE* mRNA and increased AluSz RNA; conversely, overexpression of Dicer1 increased *APOE* mRNA and decreased AluSz RNA(Supplemental Figure 5). In accordance with this line of data, knockdown of Dicer1 reduced APOE protein levels(Supplemental Figure 6). In primary mouse astrocyte cultures, Dicer1 siRNA1 increased B2 Mm2 RNA(p=0.0212) and decreased ApoE mRNA(p=0.0156) (Figures. 4A–4C). Interestingly, Dicer1 knockdown increased GFAP expression (Figure 4D). With northern blot and fluorescence *in situ* hybridization, we found that Dicer1 knockdown increased B2 Mm2 RNA (Figures. 4E and 4F). We further transfected B2 Mm2 RNA and its antisense in primary mouse astrocyte cultures and the former decreased *APOE* mRNA(p=0.0433) and APOE protein whereas its antisense had no effect on *APOE* mRNA and the protein (Figures. 4G and 4H).

**Figure 4.**
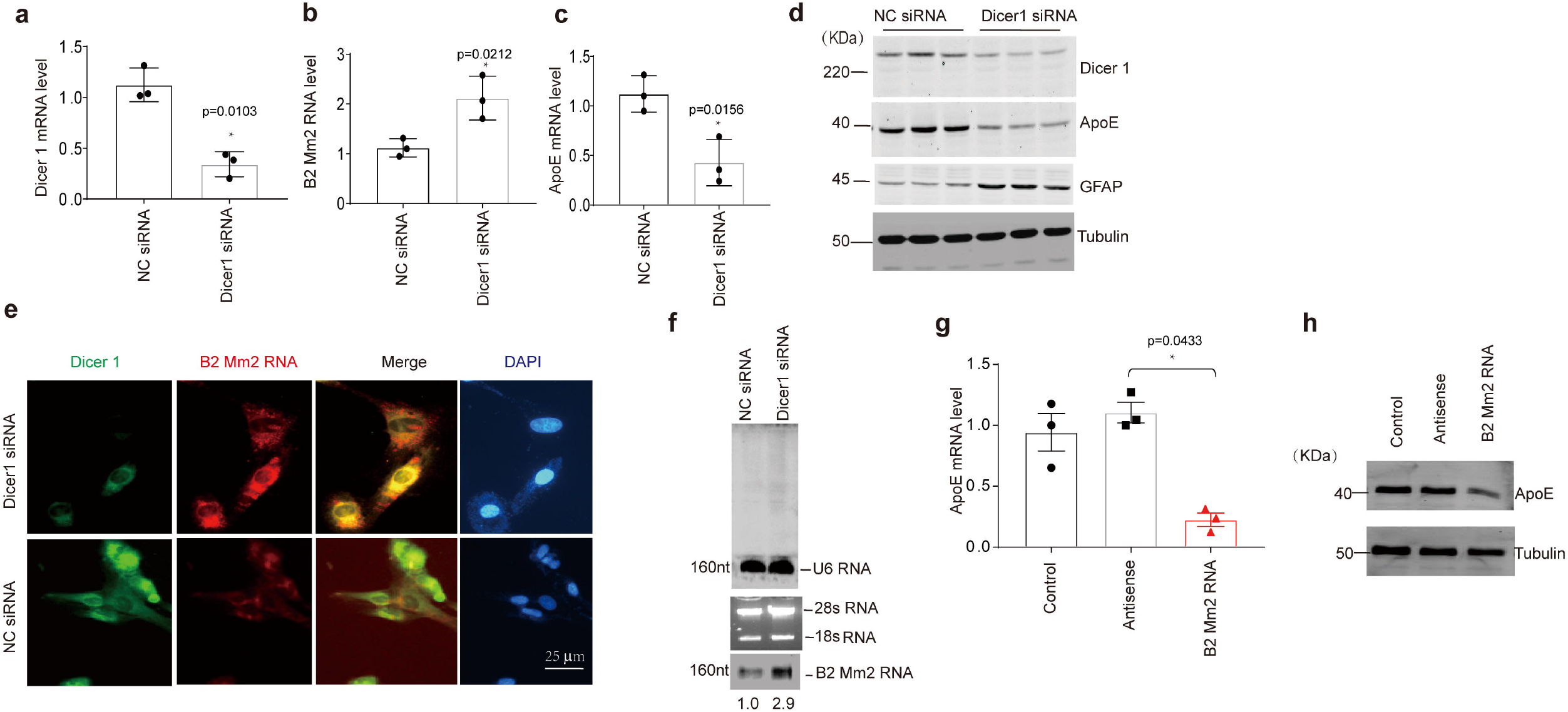
Dicer1 knockdown increased B2 Mm2 RNA and transfection of B2 Mm2 RNA increased APOE expression in primary astrocyte cultures. **(a)** Transfection of Dicer1 siRNA reduced Dicer1 mRNA levels compares with transfection of scrambled siRNA (NC siRNA) in primary astrocyte, * p=0.0103. **(b)** Transfection of Dicer1 siRNA increased B2 Mm2 RNA levels compares with transfection of scrambled siRNA (NC siRNA), * p=0.0212. **(c)** Transfection of Dicer1 siRNA reduced *APOE* mRNA levels compares with transfection of scrambled siRNA (NC siRNA), * p=0.0156. n=3. (**d**) Dicer1 was knocked down in the astrocyte cultures with Dicer1 siRNA and the protein levels of Dicer1, APOE, GFAP and Tubulin were examined with western blot. Representative image was indicated. **(e)** The primary mouse astrocyte cultures were transfected with Dicer1 siRNA for 24 hours and fixed for immunofluorescence assay. Dicer1 protein (green) was stained with anti-Dicer1 antibody and B2 Mm2 RNA (red) with fluorescence *in situ* hybridization method, respectively. DAPI was used to indicate nuclei. Scale bar, 25 μm. (**i**) Total RNA extracted from primary mouse astrocyte cultures transfected with Dicer1 siRNA and B2 Mm2 RNA was examined with Northern blot. U6 RNA was used as a RNA loading control and 28s/18s RNAs were used to indicate integrity of RNA samples. The optic density values for B2 Mm2 RNA were indicated below B2 Mm2 RNA bands. **(f)** Total RNA was extracted from hippocampus of ~11-months old WT or APP/PS1 mice. B2 Mm2 RNA was examined with Northern blot and representative image for B2 Mm2 (−160nt) RNA was indicated. **(g)** The hippocampus of APP/PS1 mice contained higher amount of B2 Mm2 RNA than WT mice at similar ages from j. Mann-Whitney *U*-test was used to compare the difference. ** p=0.0022. n=6. **(h)** Transfection of synthesized B2 Mm2 RNA reduced *APOE* mRNA. Astrocytes were plated at the density of 5X10^5^ astrocytes in 6-well plates and transfected with equal amount of synthesized B2 Mm2 or its antisense RNA at 5 nM for 48 hours. ApoE mRNA levels were examined by qRT-PCR. Paired Student’s *t-test* was used to compare the differences between two groups, * p=0.0433. n=3. **(i)** APOE protein levels in primary cultured astrocytes were reduced with transfection of synthesized B2 Mm2 RNA. The typical image was indicated. n=2.

### 3.4. B2 Mm2 RNA down-regulated mouse or human *APOE* via interacting with the Alu elements downstream of its 3’-UTR

To explore the underlying mechanism by which B2 Mm2 RNA down-regulated *APOE* mRNA, we checked the 3’-UTR of mouse *APOE* mRNA and found that an inverted Alu RNA-like sequence, B2 Mm2 RNA, located downstream of its 3’-UTR (Figure 5A). The mouse *APOE* mRNA 3’-UTR and the sequence spanning 3’-UTR and the inverted B2 Mm2 RNA were cloned into a luciferase reporter, termed pGL6-Luc-3’-UTR or pGL6-Luc-3’-UTR-B2 Mm2, respectively. Simultaneous transfection of B2 Mm2 RNA or Dicer1 siRNA decreased luciferase activity of pGL6-Luc-3’-UTR-B2 Mm2 compared with transfection of antisense RNA or scrambled siRNA(p=0.0016 for transfection of B2 Mm2 RNA and p=0.0139 for transfection of Dicer1), respectively (Figure 5B). By contrast, transfection of B2 Mm2 antisense RNA did not change luciferase relative to sham transfection (Figure 5B). Co-transfection of B2 Mm2 RNA reduced the luciferase of pGL6-Luc-3’-UTR-B2 Mm2 compared with co-transfection of pGL6-Luc-3’-UTR (p=0.0002);similarly, transfection of Dicer1 siRNA reduced the luciferase of pGL6-Luc-3’-UTR-B2 Mm2 compared with co-transfection of pGL6-Luc-3’-UTR(p=0.0002) (Figure 5C). As a control, transfection of B2 Mm2 RNA or Dicer1 siRNA did not change luciferase activity of pGL6-Luc-3’-UTR compared with sham transfection or scrambled siRNA, respectively (Figure 5C). Regarding the different effects of Dicer1 siRNA or B2 Mm2 RNA on pGL6-Luc-3’-UTR or pGL6-Luc-3’-UTR-B2 Mm2, we project that the B2 Mm2 RNA sequence downstream of *APOE* mRNA 3’-UTR was critical for this reduction. B2 RNA and its antisense RNA were synthesized as described in the materials and methods and analyzed in non-denaturing agarose gel(Supplemental Figure 7). We further checked the characteristic of human *APOE* mRNA and found three inverted tandem Alu-like sequence located downstream of its 3’-UTR. We cloned the 3’-UTR and the sequence containing the 3’-UTR plus different portions of Alu-like sequence into pGL6-Luc and termed them as pGL6-Luc-3’-UTR, pGL6-Luc-3’-UTR-AluSz, pGL6-Luc-3’-UTR-AluSz-AluJo, pGL6-Luc-3’-UTR-AluSz-2XAluJo, respectively (Figure 6A). AluSz and AluJo were synthesized as described in the materials and methods and analyzed in non-denaturing agarose gel characterized(Supplemental Figure 7). Simultaneous transfection of AluSz RNA in U87MG cells decreased luciferase activity of pGL6-Luc-3’-UTR-AluSz-2XAluJo compared with sham transfection(p=0.0001); the reduction amplitude was bigger than the effect of AluSz antisense RNA on pGL6-Luc-3’-UTR-AluSz-2XAluJo(p=0.0049) (Figure 6B). Similarly, transfection of Dicer1siRNA in U87MG cells decreased luciferase of pGL6-Luc-3’-UTR-AluSz-2XAluJo compared with transfection of scrambled siRNA(p=0.0036) (Figure 6C). When co-transfection with pGL6-Luc-3’-UTR-AluSz-2XAluJo, Dicer1 siRNA reduced luciferase to biggest effect, relative to pGL6-Luc-3’-UTR, among the three Alu elements-containing constructs (p=0.0007 *vs*.pGL6-Luc-3’-UTR-AluSz; p=0.0009 *vs*.pGL6-Luc-3’-UTR-AluSz-AluJo); the effect on pGL6-Luc-3’-UTR-AluSz was similar as on pGL6-Luc-3’-UTR-AluSz-AluJo (Figure 6D). Similar to the effect on mouse 3’-UTR construct, transfection of Dicer1 siRNA in U87MG cells did not reduce luciferase of pGL6-Luc-3’-UTR compared with scrambled siRNA (Figure 6D). These data suggest that Alu RNA downstream of human ApoE 3’-UTR was critical for the effect of Dicer1 siRNA on Alu-containing luciferase reporter. We further tested luciferase upon co-transfection of ApoE AluSz RNA which reduced the luciferase of all the constructs containing Alu elements, relative to pGL6-Luc-3’-UTR, with the biggest effect on pGL6-Luc-3’-UTR-AluSz-2XAluJo(p<0.0001 *vs*. either pGL6-Luc-3’-UTR-AluSz or pGL6-Luc-3’-UTR-AluSz-AluJo); the effect on pGL6-Luc-3’-UTR-AluSz was similar as on pGL6-Luc-3’-UTR-AluSz-AluJo (Figure 6E). Compared with the effect of AluSz RNA antisense, AluSz RNA reduced luciferase of pGL6-Luc-3’-UTR (p=0.0004) which suggest that exogenous Alu element may have marginal effect on the luciferase activity via Alu elements-independent mechanism.

**Figure 5.**
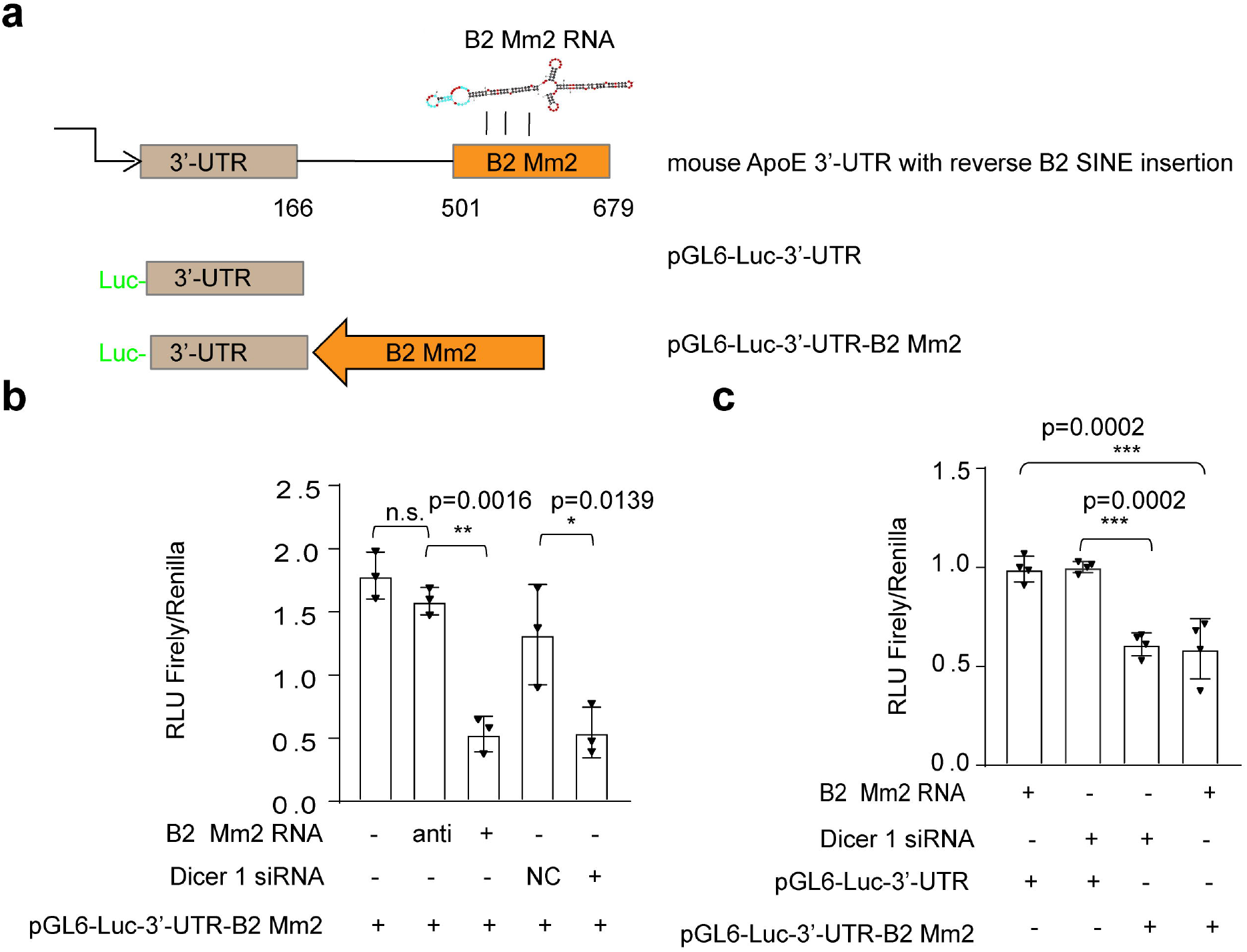
Dicer 1 knockdown or expression of B2 Mm2 RNA repressed activity of mouse *APOE* 3’-UTR-B2 Mm2 in a luciferase reporter system. **(a)** The schematic of mouse *APOE* mRNA 3’-UTR with 3’-flanking inverted B2 Mm2 SINE. The *APOE* mRNA 3’-UTR and *APOE* 3’-UTR in conjunction with inverted B2 Mm2 SINE were cloned into a pGL6-miR luciferase reporter system, termed pGL6-3’-UTR and pGL6-3’-UTR-B2 Mm2, respectively. The 3’-UTR spanned 166 bp downstream of stop codon (0 bp) and the residential inverted B2 Mm2 spanned 501bp/679bp. **(b)** The astrocytes were plated at the density of 4×104 cells/per well in 6-well plates and co-tansfected with Dicer1 siRNA or NC siRNA with pGL6-3’-UTR-B2 Mm2; or co-tranfected with synthesized B2 Mm2 RNA or its antisense with pGL6-3’-UTR-B2 Mm2 for 60 hours. The *firefly* and *renilla* luciferase activities were measured and calculated as the ratios between *firefly/renilla*, expressing as relative luminance unit (RLU). The ratios were averaged from three independent experiments. Co-transfection of Dicer1 siRNA reduced luciferase of pGL6-3’-UTR-B2 Mm2 compared with co-transfection of scrambled siRNA (NC), * p=0.0139; co-transfection of B2 Mm2 RNA decreased luciferase of pGL6-3’-UTR-B2 Mm2 compared with co-transfection of its antisense (anti), ** p=0.0016; transfection of antisense of B2 Mm2 RNA *vs*. sham transfection, n.s. n=3. One-way ANOVA followed by Tukey’s *post hoc* test was used to compare the differences. **(c)** Different luciferase activities of pGL6-3’-UTR and pGL6-3’-UTR-B2 Mm2 in response to co-transfection with Dicer1 siRNA or B2 Mm2 RNA. Upon co-transfection with Dicer1 siRNA, the luciferase activities of pGL6-3’-UTR-B2 Mm2 were decreased compared with co-transfection of pGL6-3’-UTR, *** p=0.0002; Upon cotransfection with B2 Mm2 RNA, the luciferase activities of pGL6-3’-UTR-B2 Mm2 were decreased compared with co-transfection of pGL6-3’-UTR, *** p=0.0002. n=4. One-way ANOVA followed by Tukey’s *post hoc* test was used to compare the differences among groups.

**Figure 6.**
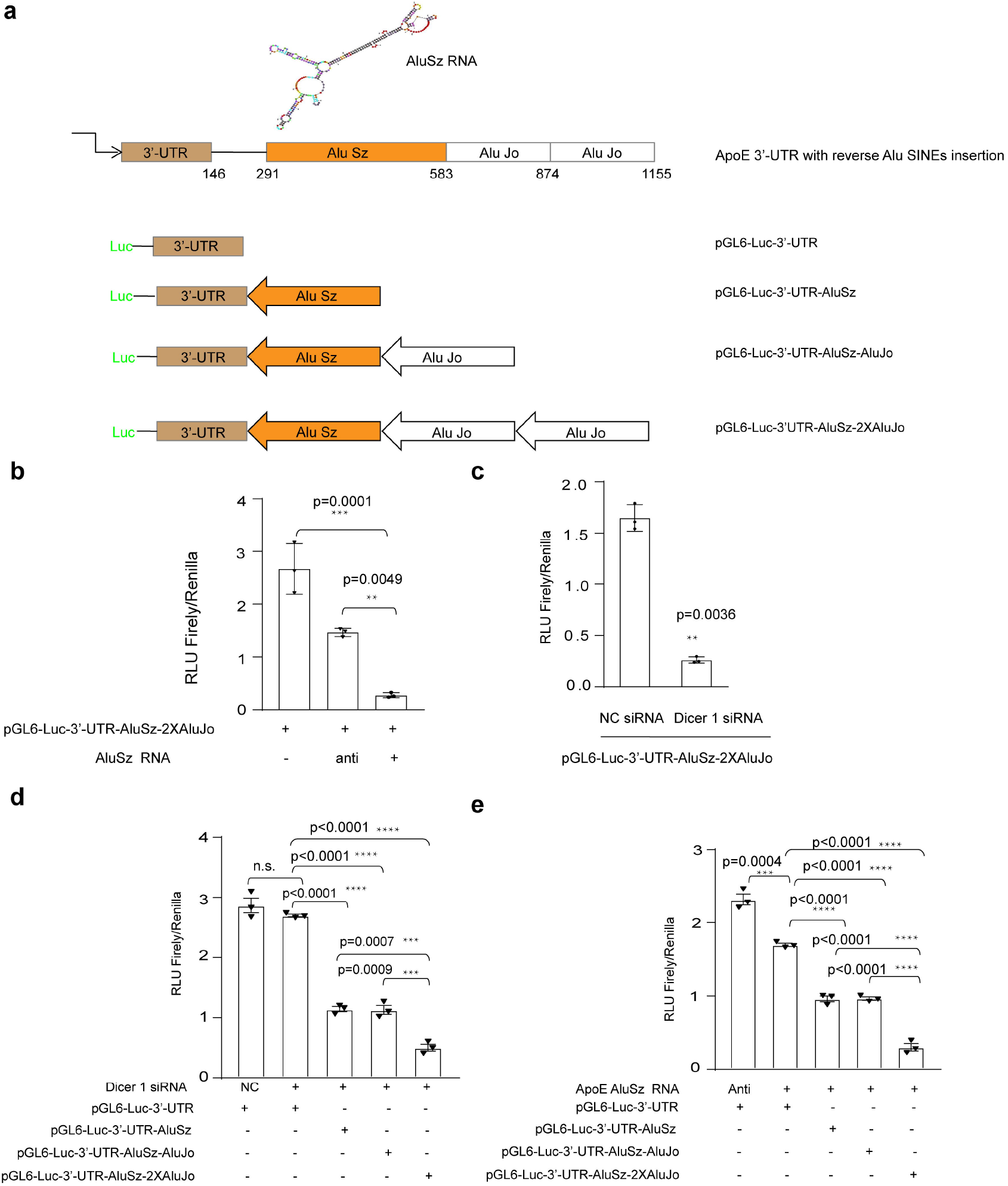
Dicer 1 knockdown or expression of AluSz RNA repressed activity of human *APOE* 3’-UTR-AluSz-2XAluJo in a luciferase reporter system. **(a)** The schematic of human *APOE* mRNA 3’-UTR with 3’-flanking tandem inverted Alu elements including AluSz SINE and two AluJo SINEs. *APOE* mRNA 3’-UTR and 3’-UTR plus different portions of Alu elements were cloned into a pGL6-miR basic luciferase reporter plasmid and termed pGL6-3’-UTR, pGL6-3’-UTR-AluSz, pGL6-3’-UTR-AluSz-AluJo, and pGL6-3’-UTR-AluSz-2xAluJo, respectively. **(b)** U87MG cells were plated at the density of 6X10^4^ cells/per well in 48-well plates until ~50% confluency and co-tansfected with synthesized B2 Mm2 RNA or B2 Mm2 RNA antisense with pGL6-3’-UTR-AluSz-2xAluJo. Co-transfection of AluSz RNA reduced the luciferase compared with sham transfection (*** p=0.0001) which indicated bigger amplitude than cotransfection of AluSz antisense(** p=0.0049). One-way ANOVA followed by Tukey’s *post hoc* test was used to compare differences among groups. n=3. (**c**) U87MG cells were plated at the density of 5X10^4^ cells/per well in 48-well plates until ~50% confluency and co-tansfected with Dicer1 siRNA or NC siRNA with pGL6-3’-UTR-AluSz-2xAluJo. Co-transfection of Dicer1 siRNA reduced the luciferase compared with transfection of scrambled siRNA. Paired Students’ *t-test* was used to compare differences. ** p=0.0036, n=3. (**d**) U87MG cells were plated at the density of 7X10^4^ cells/per well in 48-well plates until ~50% confluency and co-tansfected with Dicer1 siRNA along with pGL6-3’-UTR or Alu elements-containing luciferase reporter. Transfection of Dicer1 siRNA did not change luciferase of pGL6-3’-UTR relative to transfection of scrambled siRNA (NC) whereas Dicer1 siRNA reduced luciferase of pGL6-3’-UTR-AluSz-2xAluJo more than of pGL6-3’-UTR-AluSz (*** p=0.0007) or of pGL6-3’-UTR-AluSz-AluJo (*** p=0.0009). Transfection of Dicer1 had similar effect on pGL6-3’-UTR-AluSz as on pGL6-3’-UTR-AluSz-AluJo. Transfection of Dicer1 siRNA reduced luciferase activities of all the Alu RNA-containing constructs compared with the effect on pGL6-3’-UTR (p<0.0001 for all the comparisons). One way ANOVA followed by Tukey’s *post hoc* test was used to compare differences among groups. n=3. (**e**) U87 MG cells were plated at the density of 3X10^4^ cells/per well in 48-well plates until ~50% confluency and co-tansfected AluSz RNA or its antisense with pGL6-3’-UTR or Alu elements-containing luciferase reporter. Co-transfection of AluSz RNA reduced luciferase of pGL6-3’-UTR-AluSz-2xAluJo more than of pGL6-3’-UTR-AluSz (**** p<0.0001) or of pGL6-3’-UTR-AluSz-AluJo (**** p<0.0001). Transfection of AluSz RNA had similar effect on pGL6-3’-UTR-AluSz as on pGL6-3’-UTR-AluSz-AluJo. Transfection of AluSz RNA reduced luciferase activities of all the Alu RNA-containing constructs compared with the effect on pGL6-3’-UTR (**** p<0.0001 for all the comparisons). Transfection of AluSz RNA reduced luciferase of pGL6-3’-UTR compared with transfection of its antisense on pGL6-3’-UTR (Anti) (*** p=0.0004). One way ANOVA followed by Tukey’s *post hoc* test was used to compare differences. n=3.

## 4. Discussion

In this study, we indicated that delivery of Dicer1-expressing virus into the hippocampus improved spatial learning in ~11-month-old APP/PS1 mice. However, the extent of alleviation was lower than the effect on ~4-month-old APP/PS1 mice[24] which suggest that overexpression of Dicer1 in AD may bring more benefit in early stage of AD compared with advanced AD. The three common APOE isoforms arise from single-nucleotide polymorphisms and derives from the same gene transcript. However, the roles of them involved in AD are distinct. For example, *APOE\** ε4 increases the risk to develop AD and AD prevalence whereas *APOE\** ε2 decreases this risk and also delays the age of onset, compared with *APOE\** ε3[26, 27], a most popular *APOE* allele. Thus, we project whether delivery of Dicer1 in the central nervous system promotes amyloid clearance in humans depends on APOE isoform since APOE isoforms disrupt soluble Aβ clearance at the blood-brain barrier in an order of efficacy: ApoE4> E3> E2 [28].

Dicer1 depletion results in many defects. In embryonic stem cell, depletion of Dicer1 leads to defects in epigenetical modification of heterochromatin, resulting in inability to differentiation [29]. In post-mitotic cells, Dicer1 depletion leads to toxicity. For instance, deficiency of Dicer1 in neurons leads to neurodegeneration, resulting in microcephaly [30] whereas selective knockout of Dicer1 in astrocytes results in non-cell-autonomous neurodegeneration mediated by dysfunctional or reactive astrocytes [31]. In oligodendrocytes, deletion of Dicer1 leads to disruption of myelination, lipid accumulation and oxidative stress, resulting in a different form of non-cell-autonomous neurodegeneration [32]. The mechanisms underlying the different forms of neurodegeneration are possibly due to deficiency of mature microRNAs or accumulation of Alu RNAs. However, deletion of the component of Drosha or Argonaute2, a component of RNA-inducing silencing complex or other miRNAs-processing enzymes in RPEs, do not lead to RPE degeneration and RPE degeneration caused by Dicer1 depletion can be rescued by supplementing Alu RNA antisense [22]. These data suggest that defect of mature miRNAs syntheses in RPEs may not be a critical mechanism underlying RPE degeneration due to Dicer1 depletion. Considering that RPE is a derivative of optic neuroepithelium [33], Alu RNA accumulation rather than defect of miRNAs synthese may account for neurotoxicity in neurons under the context of Dicer1 depletion. Alu RNAs mediate cellular toxicity, either via activation of inflammasome [34] or interference with gene transcription or integration into the genome [35] [36] in which the weight of each effect accounting for neurodegeneration will be investigated in the future.

In our previous study, we have demonstrated that knockdown of Dicer1 leads to less elaborate neuronal dendrites and overexpression of Dicer1 protects against β-amyloid-induced neurotoxicity[24]. Herein, overexpression of Dicer1 exerted beneficial effects in AD mice, possibly in several aspects including alleviation of amyloid neurotoxicity, elevation of amyloid clearance, and reduction of astrogliosis. The mechanism by which Dicer1 increased amyloid clearance may result from increasing APOE expression.

Alu elements consist of two dimers derived from 7SL RNA gene, separating by a polyA linker and contain a stretch of poly A sequence in its 3’-terminus whereas at the beginning of the elements, Alu RNAs harbor a RNA polymerase III transcriptional promoter which initiates its transcription and subsequent amplification [37]. Alu elements integrate into human genome under the indispensable support of LINE-1 element, an autonomous retroelement in humans[36]. When present in 3’-UTR, Alu RNA can base-pair with cytosolic Alu RNA and long non-coding RNA to activate Staufen1-mediated mRNA decay [21]. Another evidence indicate that when present in 3’-UTR of product mRNA in the form of inverted Alu RNA, Staufen1 helps retain the mRNA in the nucleus to prevent from translation [38]. Transfection of AluSz RNA in astrocyte cells reduced *APOE* mRNA levels which suggest that Alu RNA mediates *APOE* mRNA degradation. Notably, B2 Mm RNA or tandem Alu element appears as inverted sequence downstream of *APOE* 3’-UTR. Transfection of Alu RNA or B2 MmRNA reduced the luciferase activities of the constructs suggest that cytosolic Alu RNA may collaborate with residential Alu RNA such as by base-pairing to mediate the effect. Piercing the data together, we project that exogenously applied Alu RNA or endogenous Alu RNA due to Dicer1 depletion mediates *APOE* mRNA degradation by base-pairing with Alu RNA, located downstream of 3’-UTR of *APOE* mRNA. B2 Mm2 RNA was increased in AD mouse hippocampus compared with that in WT mouse hippocampus which was in contrast with similar APOE levels between WT and AD mice. The results suggest that APOE regulation *in vivo* was more complicated than expectation. For example, APOE level is transcriptionally induced by peroxisome proliferator-activated receptor gamma (PPARgamma), liver X receptors (LXRs) in coordination with retinoid X receptors (RXRs) in which PPARgamma and LXRs have been implicated in AD therapy [39, 40].

β-Amyloid is derived from sequential cleavage of β-amyloid precursor protein by β- and γ-secretase. Apart from degradation by insulin-degrading enzyme and metalloprotease neprilysin [41, 42], β-amyloid can be uptaken, internalized, and degraded in astrocytes via low-density lipoprotein receptor-related protein 1[7] or in microglia via triggering receptor expressed on myeloid cells 2 [43]. Additionally, β-amyloid can be driven across blood-brain barrier [28] or cleaned via glymphatic system [44] during which lipidated APOE facilitates amyloid transport. In this study, we found that Dicer1 increased APOE and reduced amyloid content in the hippocampus which suggests that Dicer1 increases β-amyloid clearance via modulation of APOE. However, the effect occurred indirectly via blocking the inhibitory effect of B2 RNA on *APOE* mRNA. In addition, increasing expression of Dicer1 reduced reactive astrocytes and assumed to mitigate subsequent neuroinflammation. Altogether, overexpression of Dicer1 benefits AD mice via effects including increasing amyloid clearance and decreasing astrogliosis. Thus, our study may suggest a novel way to treat AD.

## Authors’ contributions

Wang Y conceived of the project, conducted experiments, analyzed the data, participated in writing; Lian ML conducted experiments; Song LP participated in animal husbandry; Wu S conceptualized the project, designed experiments, provided funding, analyzed the data, and wrote the manuscript.

## Acknowledgment

The project was sponsored by key projects of Wenzhou Medical University (QTJ100001) and Natural Science Foundation of Zhejiang Province (LY18H120003).

