## Supplemental Figures and legends for "Dicer1 promotes Aβ clearance via blocking B2 RNA-mediated repression of apolipoprotein E"

### Supplemental figures and figure legends

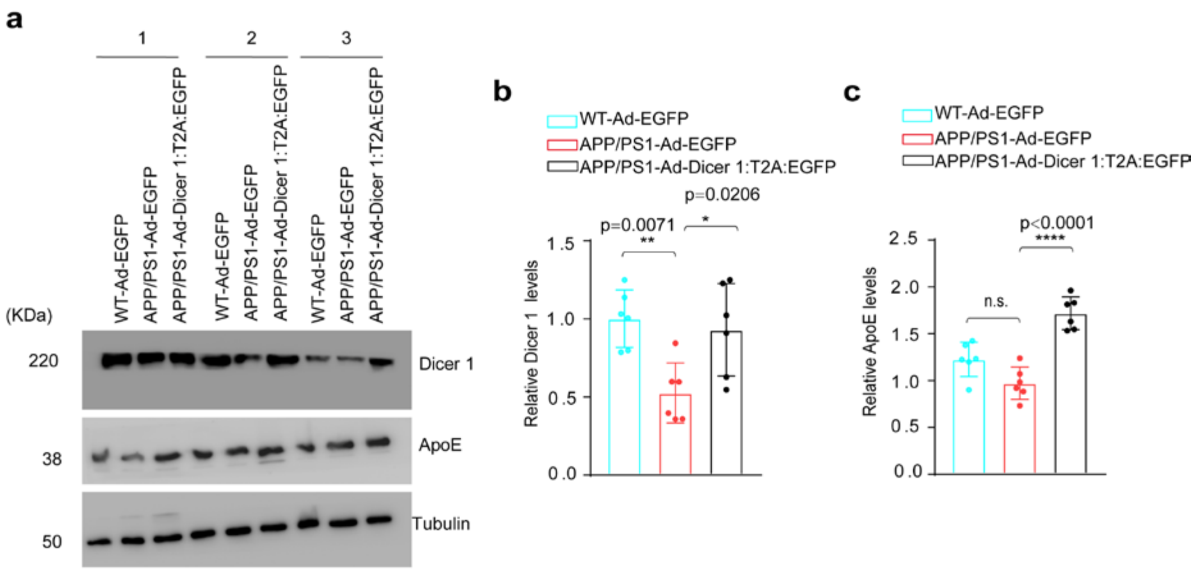


**Figure S1**. **Overexpression of Dicer 1 induced APOE expression in the hippocampus of 4-month-old APP/PS1 mice.** **(a)** Representative blots of Dicer 1 and APOE in the hippocampus of littermate WT mice injected with vehicle virus (WT-Ad-EGFP), APP/PS1 mice injected with vehicle virus (APP/PS1-Ad-EGFP), APP/PS1 mice injected with Dicer1-expressing virus (APP/PS1-Ad-Dicer1:T2A:EGFP). **(b)** Relative Dicer1 protein levels in the three groups of mice. **(c)** Relative APOE protein levels in the three groups of mice. One way ANOVA followed by Tukey's *post hoc* test was used to compare differences among groups, n=6 per group.

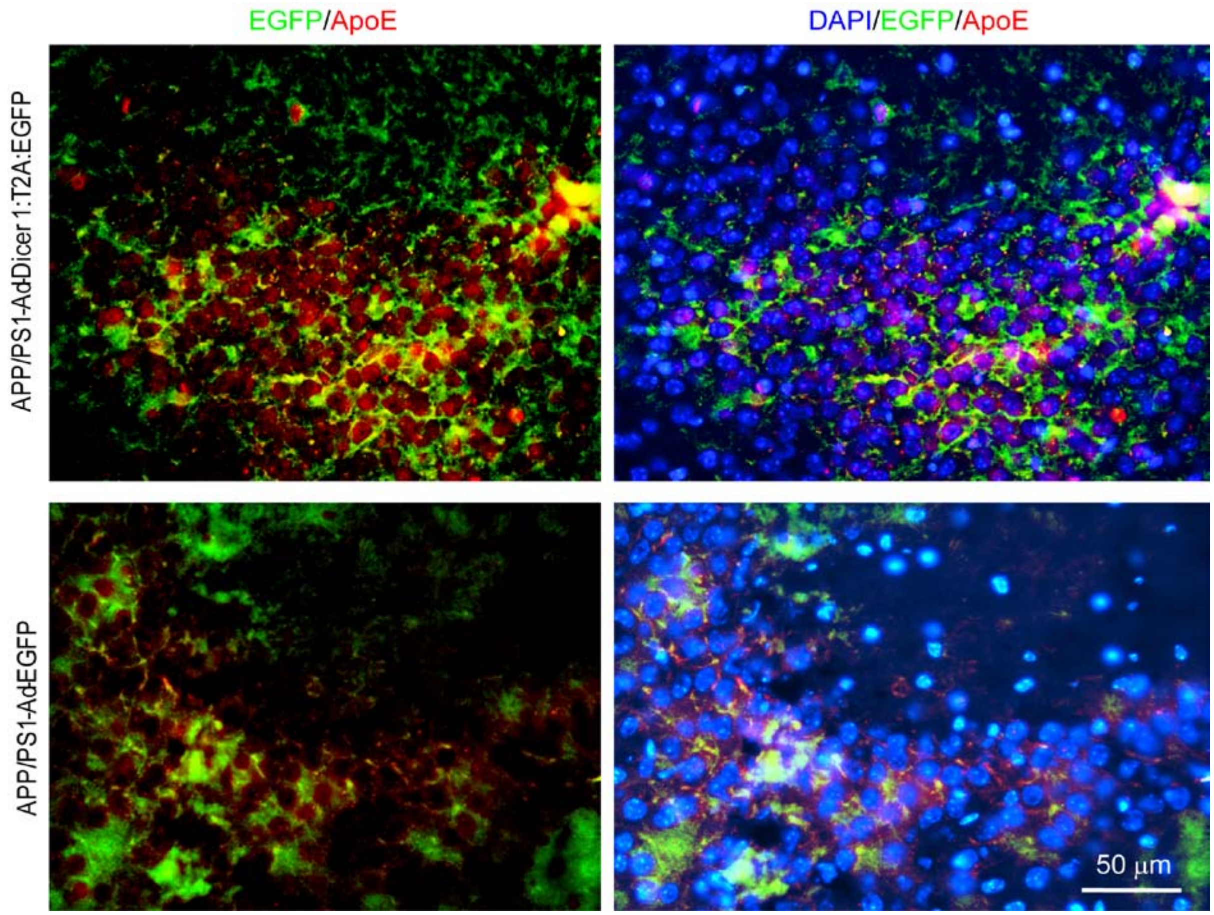


**Figure S2. Injection of Dicer1-expressing virus increased APOE.** The indicated images show immunofluorescence of APOE expression (red) and virus infection (green) in the CA3 region of hippocampus from 11-month-old APP/PS1 mice injected with Dicer1-expressing virus (top panel) or vehicle virus (low panel). DAPI was used to indicate nuclei in the images at the right column. Scale bar, 50 μm. n=2.

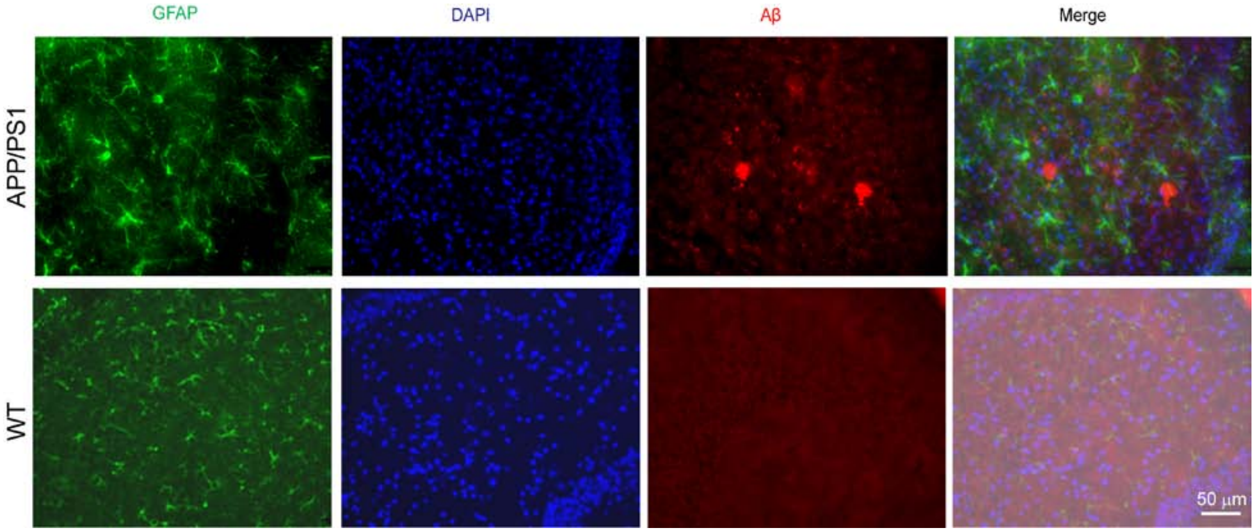


**Figure S3. Astrogliosis in APP/PS1 mice.** The representative images indicated Aβ staining (red) and astrocytes indicated by GFAP staining (green) in cerebral cortical cryosections from 11-month-old APP/PS1 mice (top panel) or WT mice with similar age (low panel). DAPI was used to indicate nuclei and all the staining were merged at the fourth column. Scale bar, 100 μm. n=2.

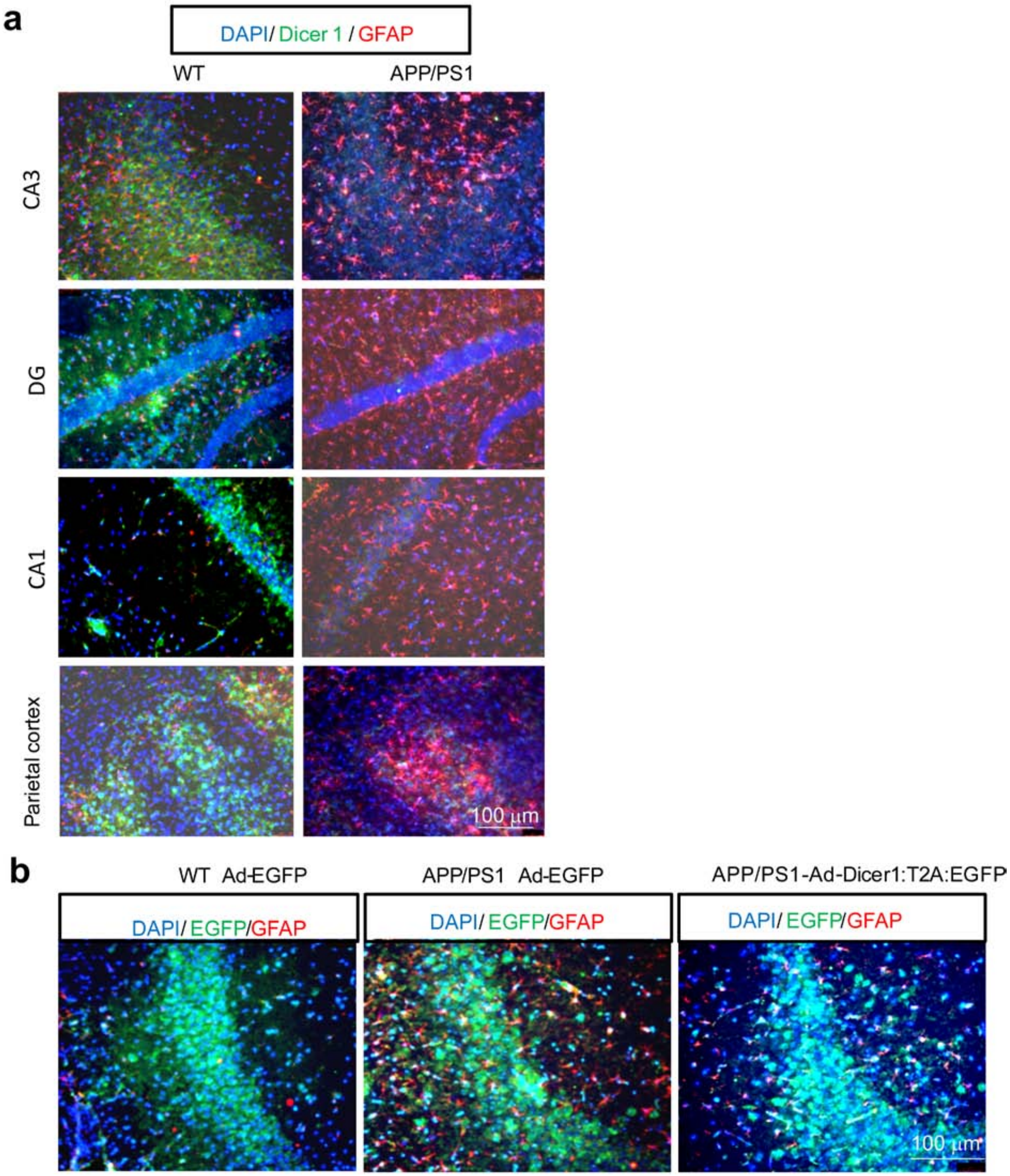


**Figure S4. Intrahippocampal injection of Dicer1-expressing virus reduced the amounts of reactive astrocytes in the hippocampus of 11-month-old APP/PS1**

**mice. (a)**The brain sections of 11-month-old APP/PS1 mice or WT mice were double-stained with Dicer1 antibody (green) and GFAP antibody (red). The representative images of CA1, CA3 and DG regions of hippocampus and parietal cortex were shown, respectively. Scale bar, 100 μm. n=3. **(b)** WT or APP/PS1 mice at 11 months were injected with 2 µL of Ad-pCMV-EGFP or Ad-pCMV-Dicer1:T2A:EGFPvirus ($1.2 \times 10^9$ vg/mL) into the hippocampus as described in the materials and methods. The brain sections from each group were stained with anti-GFAP antibody (red). DAPI was used as nuclei staining for all sections and the merged images of DAPI/EGFP/GFAP in the CA3 region were shown. Scale bar, 100 μm. n=3.

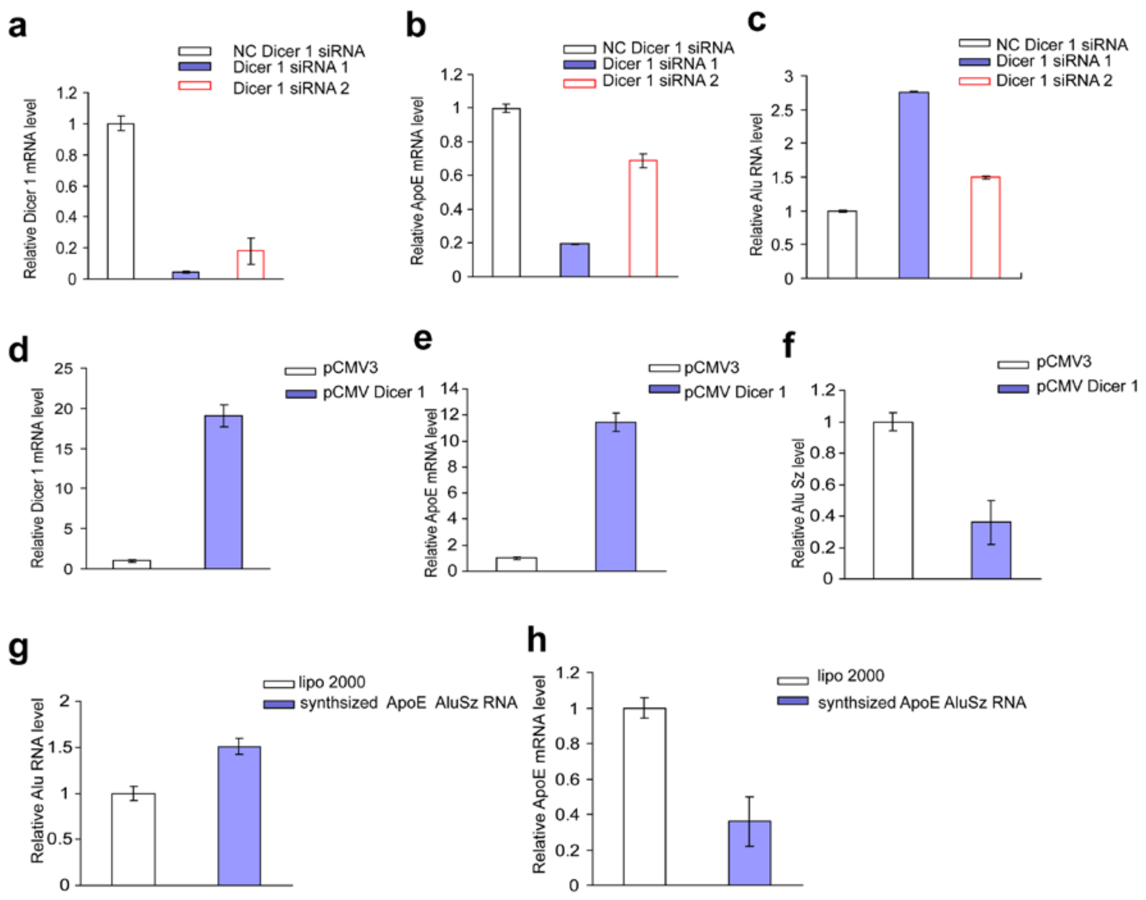


**Figure S5. The effects of Dicer1 knockdown/overexpression on the levels of *APOE* mRNA and AluSz RNA in U87 MG cells which were determined by quantitative reverse-transcriptase PCR (qRT-PCR).** The plated U87 MG cells were transfected with NC siRNA, Dicer1 siRNA 1 or Dicer1 siRNA2 for 60 hours, respectively. **(a)** Relative amount of Dicer1 mRNA level under indicated conditions. **(b)**Relative amount of *APOE* mRNA level under indicated conditions. **(c)** Relative amount of AluSz RNA level under indicated conditions. The plated U87MG cells were transfected with empty vector, pCMV3 or Dicer1-expressing plasmid, pCMV Dicer1 for 60 hours. **(d)**Relative amount of Dicer1 mRNA level under indicated conditions. **(e)** Relative amount of *APOE* mRNA level under indicated conditions. **(f)** Relative amount of AluSz RNA level under indicated conditions. The plated U87 MG cells were also transfected with lipofectamine 2000 (sham transfection), or synthesized AluSz RNA (7 µg) for 60 hours. (**g**) Relative amount of Alusz RNA level under indicated conditions. (**h**) Relative amount of *APOE* mRNA level. n=2 cell culture preparations for the experiments in a-h.

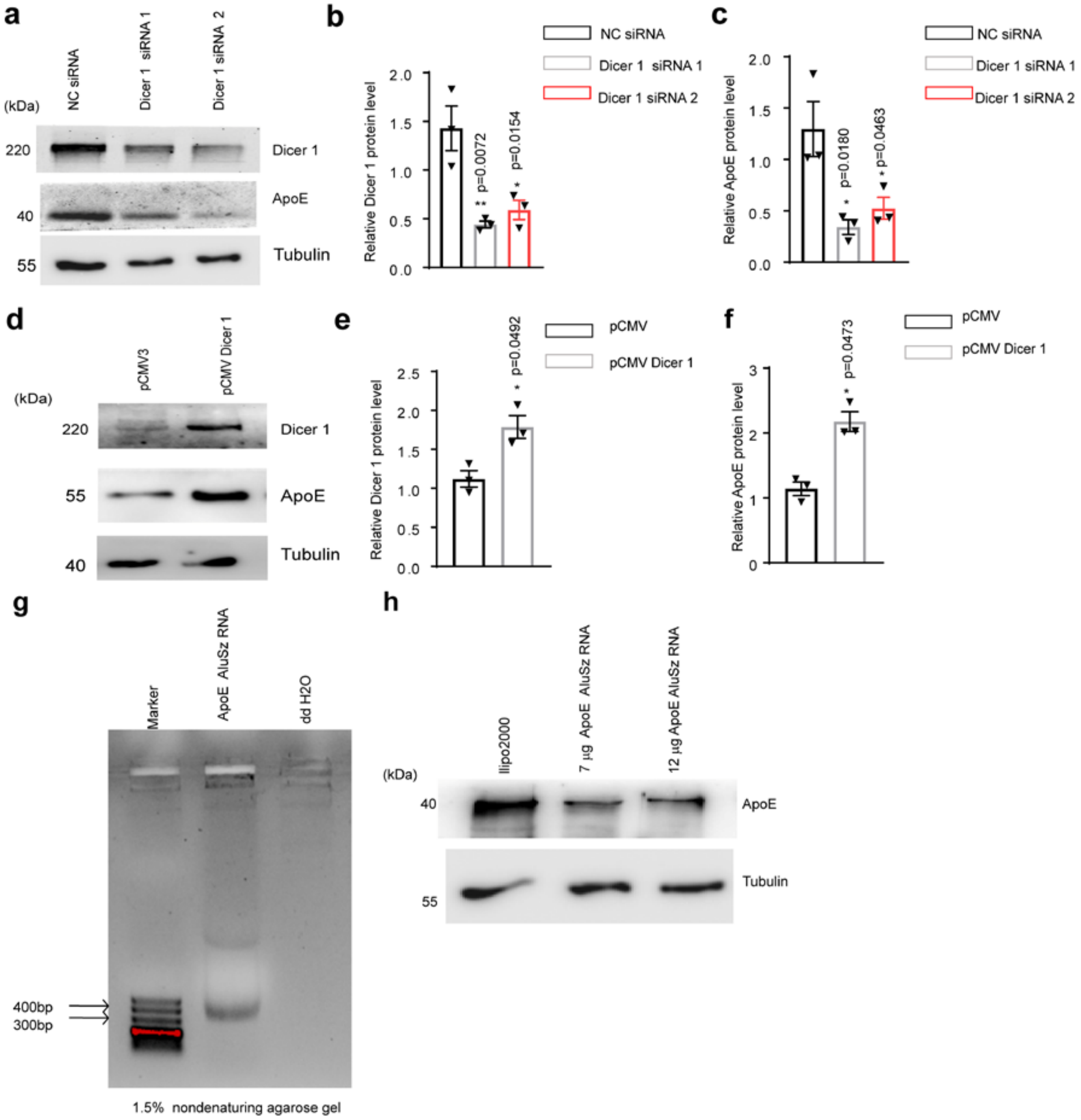


**Figure S6. The effect of Dicer1 knockdown/overexpression on the protein levels of APOE. (a)** The plated U87MG cells were transfected with scrambled siRNA (NC siRNA) and two sets of Dicer1 siRNA. Dicer1and APOE protein levels were examined with western blot. Tubulin was used as an internal loading control. n=3. **(b)** Relative Dicer1 protein levels under indicated conditions. **(c)** Relative APOE protein levels under indicated conditions. **(d)** The plated U87MG cells were transfected with empty vector, pCMV or Dicer1-expressing vector, pCMV Dicer1 plasmid, for 60 hours. Dicer1 and APOE protein levels were examined with western blot and Tubulin used as an internal loading control. n=3. **(e)** Relative amount of Dicer1 protein under indicated conditions. **(f)** Relative amount of APOE protein under indicated conditions. **(g)** The synthesized AluSz RNA was examined on 1.5 X non-denaturing agarose gel and the bands was located among ~300bp~400bp. **(h)** Relative amount of APOE protein under indicated conditions. The plated U87MG cells were transfected with lipofectamine 2000 (sham transfection), or transfected with 7-/12-µg synthesized *APOE* AluSz RNA for 60 hours. n=2 cell culture preparations for all the experiments in a-h.

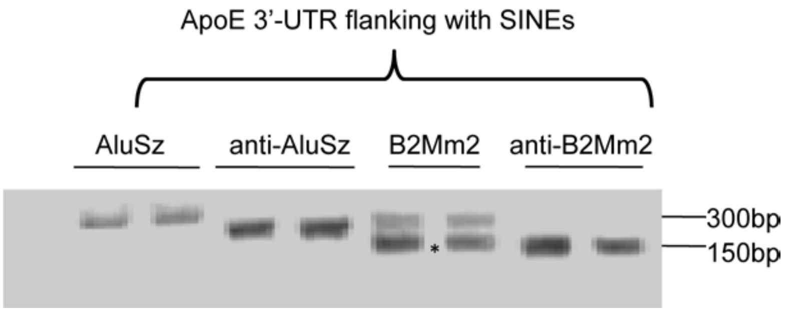


**Figure S7. The synthesized Alu RNAs were resolved on 1.5% non-denaturing agarose gel.** The human *APOE* AluSz SINEs and its antisense RNA, the mouse *APOE* B2 Mm2 RNA and its antisense were synthesized as described in the materials and methods and analyzed in 1.5% non-denaturing agarose gel.
